# Trigeminal Nerve Stimulation Attenuates Traumatic Brain Injury–Induced Neuronal Damage via CGRP-PINK1 Mitophagy Axis

**DOI:** 10.64898/2026.10.12.758678

**Authors:** Chunyan Li, Keren Powell, Steven Wadolowski, Junhwan Kim, Kaitlyn Rams, Bernadette Tam, Amir Dehdashti, Jamie S. Ullman, David LeDoux, Lance B. Becker, Ping Wang

## Abstract

Secondary damage after traumatic brain injury (TBI) is driven by progressive mitochondrial failure, yet clinically viable strategies to directly safeguard mitochondrial networks remain a major therapeutic gap. Here, we show that non-invasive trigeminal nerve stimulation (TNS), which engages the brain’s endogenous oxygen-conserving diving reflex, preserves neuronal survival by directly regulating the mitochondrial PINK1/Parkin quality-control pathway. We demonstrate that TBI induces the pathological aggregation of the outer mitochondrial membrane receptor TOM20, in a CGRP-dependent manner, which affects binding of the damage-sensor PINK1 and renders downstream Parkin-mediated rescue ineffective. TNS-induced neuronal CGRP release prevents this TOM20 aggregation, restoring proper PINK1 function and downstream Parkin activation. This preservation of mitochondrial architecture reinforces the outer membrane’s barrier function, limiting pro-apoptotic cytochrome c leakage and supporting NRF2 antioxidant defenses, ultimately reducing progressive infarct expansion and recovering sensorimotor functions. The significance of this work lies in bridging physical neuromodulation directly to the preservation of organelle-level architecture. By demonstrating that stimulation of trigeminal afferents to recruit an oxygen-conserving reflex can stabilize outer membrane signaling architecture and preserve mitochondrial survival machinery, this study outlines a non-pharmacological strategy to protect mitochondrial function and inhibit acute post-traumatic neurodegeneration.

## 1. Introduction

Traumatic brain injury (TBI) initiates a delayed and evolving secondary injury response that contributes substantially to neurological disability [1–3]. Following the initial mechanical insult, excitotoxicity, oxidative stress, neuroinflammation, blood–brain barrier dysfunction, and mitochondrial failure interact to promote neuronal injury [4–6]. Despite advances in neurocritical care, no approved therapy specifically interrupts these interconnected secondary-injury processes [3]. Mitochondria are central to this cascade because impaired oxidative phosphorylation, ATP depletion, calcium overload, and excessive reactive oxygen species (ROS) production compromise neuronal bioenergetics and activate cell-death pathways [7–9]. Furthermore, damaged mitochondria release factors that act as danger signals, amplifying innate immune activation and tissue injury. These observations identify mitochondrial quality control as an important regulator of outcome after TBI and a potentially actionable therapeutic target.

Mitophagy, the selective elimination of damaged mitochondria through autophagic– lysosomal degradation, is a central component of mitochondrial quality control and contributes to neuronal homeostasis under stress [10,11]. By removing dysfunctional mitochondria, effective mitophagy can limit ROS accumulation, preserve bioenergetic capacity, reduce mitochondrial danger signaling, and restrain inflammatory and cell-death pathways [11]. Experimental studies indicate that TBI alters mitochondrial turnover and mitophagy-related signaling during the early post-injury period and that impaired mitochondrial clearance can influence neuronal injury and behavioral outcome [12–14]. However, the effects of mitophagy following TBI are highly context-dependent and may vary according to timing, magnitude, and cellular compartment [15,16]. Therefore, an effective therapeutic strategy is not simply to elevate mitophagy-associated proteins, but to resolve pathway dysregulation, prevent further mitochondrial structural damage, and reacquire functional integrity within injured neurons.

Trigeminal nerve stimulation (TNS) is a noninvasive neuromodulation strategy with emerging neuroprotective potential after TBI [17–21]. While some applications of TNS have been shown to improve cerebral perfusion, reduce edema, and attenuate inflammation, a mechanistically distinct approach utilizes precisely tuned TNS to engage the mammalian diving reflex, a potent endogenous oxygen-conserving program [22–24]. Our previous work established that diving reflex application confers robust neuroprotection [25,26], an effect mediated by significant antioxidant response upregulation and the activation of the key cytoprotective transcription factor, nuclear factor erythroid 2-related factor 2 (NRF2) [25]. Crucially, NRF2 is not only a key regulator of antioxidant defenses [27] but also functions as an upstream controller of mitochondrial homeostasis and quality control [28]. This established link provides a direct mechanistic bridge between our TNS paradigm and the potential for preserving neuronal viability by stabilizing downstream mitochondrial pathways after TBI.

The molecular mediator linking TNS to this protective cascade is calcitonin gene-related peptide (CGRP). As a principal neuropeptide of the trigeminal system released during TNS [29], CGRP modulates cellular responses to injury and stress [30–32], and our prior research confirmed that CGRP release is critical for NRF2 activation during oxygen-conserving diving reflex [25]. Crucially, NRF2 signaling is known to intersect with direct mitochondrial quality control pathways; for instance, NRF2 activation can promote mitophagy and has been shown to partially reverse neurodegeneration associated with PINK1 or Parkin deficiency [33–35]. The primary pathway for the selective removal of damaged mitochondria is the PINK1/Parkin pathway [36], which becomes dysregulated and inconsistent following TBI [15,16,37–40]. In conjunction with the recognized capacity to regulate oxidative stress [30], which prevents PINK1/Parkin overload and breakdown [41], this raises an essential question: if TNS-CGRP signaling engages NRF2 and potent antioxidative responses, does it also resolve the post-traumatic dysregulation of the direct, organelle-specific PINK1/Parkin machinery? We therefore formed the central hypothesis that TNS-induced CGRP signaling preserves mitochondrial integrity after TBI by stabilizing the function of the PINK1/Parkin pathway.

To test this central hypothesis, in this study, we investigated whether tuned TNS, by engaging the brain’s endogenous oxygen-conserving diving reflex, promotes neuroprotection after TBI through regulation of the mitochondrial PINK1/Parkin quality-control pathway. We specifically hypothesized that TNS-induced CGRP release preserves the functional integrity of this pathway by stabilizing its core signaling components and maintaining the structural arrangements required for its activation. To test this, we examined key steps in the pathway, including the stability of PINK1 (PTEN-induced kinase 1) and the operational integrity of its outer membrane docking environment, which includes the import receptor TOM20 (Translocase of Outer Mitochondrial Membrane 20) [42]. We further tested whether this preservation of mitochondrial pathway function would prevent cytochrome c leakage and preserve the functional capacity of the NRF2 antioxidant system to limit secondary neuronal injury. By utilizing pharmacological CGRP inhibition, we sought to determine the direct functional dependency of these organellar, cellular, and neurological outcomes on trigeminal-CGRP signaling. This approach was designed to establish a clear mechanism linking TNS to the preservation of core mitochondrial machinery after TBI.

## 2. Materials and Methods

### 2.1 Ethical approvals, animal husbandry, and experimental design

All experimental procedures were approved by the Institutional Animal Care and Use Committee (IACUC) of the Feinstein Institutes for Medical Research under a pre-approved protocol and conducted in strict accordance with the National Institutes of Health (NIH) Guide for the Care and Use of Laboratory Animals. Male Sprague-Dawley rats (10–12 weeks old, weighing 350– 450 g; Charles River Laboratories, Kingston, NY, USA) were housed in a temperature-controlled facility under a 12-hour light/dark cycle with ad libitum access to food and water, in cages lined with Enrich-o’Cobbs bedding (The Andersons, Inc., Maumee, OH, USA). To systematically evaluate the therapeutic effects of tuned TNS and the mechanistic role of its downstream signaling mediator, CGRP, a total of 221 rats (216 after exclusion of 5 due to failure to meet baseline neurobehavioral performance criteria) were randomized across two distinct experimental protocols. In *Experiment 1*, designed to analyze the regional and temporal impact of tuned TNS following controlled cortical impact (CCI) injury, 144 animals were allocated to a 4×3 factorial design comprising four survival timepoints (1, 4, 24, and 72 hours) and three treatment conditions: sham surgery, CCI alone, and CCI with tuned TNS. Each group within this experiment (n=12 per cohort) was split equally (n=6 each) between fresh tissue collection for biochemical analysis and transcardial perfusion-fixation for immunohistochemical and histological evaluation. In *Experiment 2*, designed to determine CGRP’s role within the assessed mitophagy pathways, 72 animals were allocated across six treatment paradigms to assess the hyper-acute (4 hours) and sub-chronic (72 hours) consequences of pharmacological CGRP receptor blockade in CCI itself. These animals were distributed among sham surgery, CCI with vehicle, and CCI with the administration of the selective CGRP receptor antagonist CGRP_8-37_ (Tocris Biosciences, Bristol, UK), with each cohort (n=12) split equally (n=6 each) between fresh tissue harvesting and transcardial perfusion collection schemas. In this experiment, sham and vehicle animals received a bolus of sterile saline for parity.

### 2.2 Controlled cortical impact model of TBI and therapeutic interventions

Following the quarantine period and baseline behavioral testing, severe TBI was induced using a highly reproducible, left-sided controlled cortical impact (CCI) paradigm [18,43]. Animals were initially anesthetized with 5% isoflurane in medical-grade air and secured in a stereotaxic frame, where anesthesia was maintained at a surgical plane with 1.5% isoflurane. Core body temperature was continuously monitored via a rectal probe and maintained at a physiological 37.0± 0.2°C utilizing a homeothermic warming system. A longitudinal midline scalp incision was made, and a 6-mm diameter craniotomy was performed over the left parietal cortex, centered 4 mm lateral to the sagittal suture and equidistant between bregma and lambda, with careful preservation of the underlying dura. Cortical injury was delivered using an electromagnetic impactor (Impact One™ Stereotaxic CCI Instrument, Leica Biosystems, Wetzlar, Germany) using the following parameters: a 5-mm flat-tipped impactor tip, an impact velocity of 6.0 m/s, a deformation depth of 3.0 mm, and a dwell time of 100 ms. Sham-injured animals underwent identical surgical preparation and craniotomy but did not receive the cortical impact. Starting 30 minutes following the injury or sham procedure, therapeutic interventions were initiated; animals assigned to the tuned TNS cohort received a continuous 30-min session of tuned electrical TNS, whereas those in the pharmacological cohorts received an intravenous (IV) bolus of the CGRP receptor antagonist CGRP_8-37_ (200 μg/kg) via the tail vein. To maintain strict experimental parity across all groups, sham and vehicle-treated TBI animals in *Experiment 2* received an equivalent volume (0.1 mL) of sterile saline IV. One-hour post-injury, following the completion of the acute stimulation period, the scalp wounds of animals allocated to extended survival cohorts were closed using 4-0 silk sutures and rapid-seal wound gel. Postoperative care included subcutaneous administration of buprenorphine for systemic analgesia and topical antibiotic prophylaxis. Animals were monitored in temperature-controlled recovery chambers until they achieved sternal recumbency before being returned to their home cages.

### 2.3 Tuned non-invasive trigeminal nerve stimulation and diving reflex activation

Non-invasive tuned TNS was administered 30 minutes post-injury using a custom-engineered nasal clip stimulator designed to bilaterally target the infraorbital branch of the trigeminal nerve at the whisker pads, using previously established parameters to mimic the sensory afferent input that triggers the endogenous, oxygen-conserving diving reflex [22]. The stimulating clip apparatus utilized two physiological-grade conductive interface pads positioned bilaterally over the infraorbital nerves. Electrical stimulation was generated using an Isolated Pulse Stimulator (Model 2100, A-M Systems, Sequim, WA, USA) programmed with highly calibrated, tuned parameters: rectangular biphasic pulses delivered at a frequency of 50 Hz, with a 0.5 ms pulse width and a 2 V output amplitude. To elicit the sustained, protective physiological effects of the diving reflex without causing habituation or sensory adaptation, stimulation was applied in 20-second trains every 5 minutes over a total therapeutic session duration of 30 minutes, initiated 30 minutes following the CCI.

### 2.4 Multi-domain neurobehavioral and functional assessments

To comprehensively evaluate neurological recovery, a multi-domain neurobehavioral testing battery was performed at 72 hours post-TBI, with baseline testing conducted 24 hours prior to injury induction to establish pre-injury functional status. All assessments were performed during the light cycle by an experienced technician blinded to the treatment groups. Animal movement and exploratory behavior were tracked using EthoVision XT video tracking software (Version 17, Noldus Information Technology, Wageningen, Netherlands) and manually verified for absolute accuracy. To eliminate confounding olfactory cues, all testing apparatuses, beams, and enclosures were thoroughly sanitized with a hydrogen peroxide solution followed by 75% ethanol between animal trials. As noted in the experimental design, five animals exhibiting baseline neurological deficits or memory impairment during pre-injury screening were excluded from the study.

#### Adjusted Neurological Severity Score

Global and domain-specific sensorimotor deficits were quantified using the validated Adjusted Neurological Severity Score (A-NSS), a comprehensive composite battery adapted from traditional neurological severity scoring protocols [44]. This scale assesses motor function (including locomotor symmetry and limb flexion), sensory responses (proprioception, tactile, and visual reflexes), vestibulomotor coordination, and cranial nerve integrity. Points are assigned for the absence of physiological reflexes or the presence of abnormal behavior, with individual tasks graded on a progressive scale of impairment. The cumulative A-NSS score ranges from 0 to 45, where a score of 0 represents normal neurological function and 45 indicates maximal, severe neurological deficit.

#### Balance Beam Test

To specifically isolate vestibulomotor coordination and static equilibrium, a balance beam test was performed. Animals were placed on an elevated, narrow wooden beam (1.5 cm in width) and evaluated on their ability to maintain a stable, four-limb posture atop the beam. Performance was scored on a standardized 1-to-6 scale based on the number of limbs remaining on the beam and the duration and nature of clinging behavior. In this scoring paradigm, a score of 1 represents perfect balance, while a score of 6 indicates immediate falling or an inability to cling to the beam for at least 20-seconds, reflecting severe impairment.

#### Accelerating Rotarod Test

Dynamic motor coordination, endurance, and motor learning were assessed using an accelerating rotarod system (Rotamex 4/8, Columbus Instruments, Columbus, OH, USA) [44]. Prior to surgical intervention, all animals underwent a rigorous training protocol consisting of three daily training sessions across three consecutive days to establish a stable performance plateau. During training and testing, rats were placed on the spindle, which accelerated linearly from 4 to 30 rpm over a 300-second trial. At 72 hours post-injury, the latency to fall from the rotating rod was recorded across three consecutive trials per animal. The longest (best) latency to fall achieved by each rat was used for statistical analysis, with a maximum cutoff latency of 300 seconds assigned to animals that completed the entire trial without falling, and a training threshold of 120-seconds for inclusion.

#### Randall-Selitto Paw Pressure Test

To evaluate post-traumatic hyperalgesia and somatosensory processing, mechanical nociceptive withdrawal thresholds were determined using an electronic Randall-Selitto paw pressure test (IITC 2500 Digital Paw Pressure Meter, IITC Life Science, Woodland Hills, CA, USA) [45,46]. Each rat was gently wrapped in a soft towel and held by an investigator who had been previously familiarized with the animal to minimize stress-induced analgesia. A second, blinded investigator applied a progressively increasing mechanical force via a blunt-tipped stylus to the plantar surface of the contralateral (right) forepaw. The application of force was immediately terminated upon the presentation of a clear nociceptive response, such as paw withdrawal or vocalization, and the mechanical threshold was automatically recorded in grams.

### 2.5 Fresh tissue collection and regional cortical dissection

At the designated survival endpoints (1, 4, 24, or 72 hours post-injury), animals allocated for biochemical analysis were deeply anesthetized with isoflurane and rapidly decapitated. The brains were quickly extracted and placed on an ice-cold dissection matrix. The brain was microdissected to isolate three distinct regional tissue samples: the primary necrotic core (infarct), the salvageable penumbral region (peri-infarct cortex), and the presumed homotypic uninjured region of the contralateral cortex. These dissected cortical tissues were immediately flash-frozen in liquid nitrogen to preserve molecular integrity and stored at −80°C until further subcellular fractionation, Western blotting, or biochemical analyses.

### 2.6 Transcardial perfusion, cryoprotection, and tissue sectioning

At the designated survival endpoints, a separate cohort of animals (n = 6 per group) was deeply anesthetized and subjected to transcardial perfusion. The thoracic cavity was opened, and the left ventricle was cannulated to deliver 0.1 M ice-cold phosphate-buffered saline (PBS; Sigma-Aldrich, St. Louis, MO, USA) to clear the vasculature, followed by the infusion of 4% paraformaldehyde (PFA; Santa Cruz Biotechnology, Dallas, TX, USA) in 0.1 M PBS for in situ tissue fixation. Following perfusion, the brains were rapidly and carefully extracted, post-fixed in 4% PFA overnight at 4°C, and subsequently dehydrated through immersion in graded sucrose solutions (10%, 20%, and 30% in PBS). The cryoprotected tissues were then equilibrated and frozen in a 1:1 mixture of 30% sucrose and optimal cutting temperature (OCT) compound (Electron Microscopy Sciences, Hatfield, PA, USA) and stored at −80°C. Serial coronal sections were cut at a thickness of 14 µm using a cryostat (Leica Biosystems, Deer Park, IL, USA) at 400-µm intervals throughout the entire rostrocaudal extent of the brain, yielding 28 to 32 representative sections per specimen. The sections were mounted onto Polysine glass slides (Thermo Fisher Scientific, Waltham, MA, USA) and stored at −80°C for downstream histological and immunofluorescence staining.

### 2.7 Subcellular fractionation and Western blot analysis

Frozen cortical tissue samples were pulverized under liquid nitrogen and aliquoted for subsequent molecular analyses. To evaluate compartment-specific protein expression, mitochondrial fractions were isolated using a specialized commercial mitochondrial isolation kit acceptable for use with frozen tissues (**Supplementary Table 1**), with the explicit caveat that these frozen-extracted organelles were utilized solely for Western blot profiling and not for functional enzymatic assays. Briefly, tissues underwent repetitive mechanical homogenization and differential centrifugation in an ice-cold mitochondrial extraction buffer to isolate the pure mitochondrial pellet. To assess the nuclear translocation of NRF2, separate cytoplasmic and nuclear fractions were prepared using a commercial nuclear and cytoplasmic extraction kit (**Supplementary Table 1**). Pulverized tissues were homogenized in a cytoplasmic pre-extraction buffer and centrifuged to yield the cytoplasmic supernatant, after which the remaining nuclear pellet was resuspended in nuclear extraction buffer, subjected to serial vortexing, and centrifuged to isolate the nuclear extract. In parallel, whole-tissue lysates were prepared by homogenizing pulverized cortex in radioimmunoprecipitation assay (RIPA) lysis buffer (Thermo Fisher Scientific, Waltham, MA, USA) supplemented with a protease and phosphatase inhibitor cocktail (Thermo Fisher Scientific). Protein concentrations across all lysates, whole-tissue, cytoplasmic, nuclear, and mitochondrial—were determined and standardized using a bicinchoninic acid (BCA) protein assay. Equal amounts of protein were resolved by SDS-PAGE and transferred via semi-dry methodology using the Bio-Rad Turbo Transfer setup onto polyvinylidene difluoride (PVDF) membranes. The membranes were blocked for 1 hour at room temperature with either 5% non-fat skim milk (Thermo Fisher Scientific) or 5% bovine serum albumin (BSA; MilliporeSigma, Burlington, MA, USA) in Tris-buffered saline containing 0.1%

Tween-20 (TBST) and then incubated overnight at 4°C with validated primary antibodies (**Supplementary Table 2**). Following three washes in TBST, membranes were incubated with horseradish peroxidase (HRP)-conjugated secondary antibodies (Abcam, Waltham, MA, USA) in 5% milk for 1 hour at room temperature. Protein bands were visualized using enhanced chemiluminescence (ECL) substrate (Thermo Fisher Scientific) on a Bio-Rad ChemiDoc Imaging System. Densitometric quantification was performed using ImageJ software (NIH, Bethesda, MD, USA), with any adjustments to image brightness or contrast applied uniformly across the entire blot. To ensure compartment-specific loading accuracy, target protein signals were normalized to β-actin (for total and cytoplasmic proteins), voltage-dependent anion channel 1 (VDAC1; for mitochondrial proteins), or Lamin B1 (for nuclear proteins). Samples in Western blot experiments each underwent independent replication twice to confirm the validity of the molecular trends, and data are presented as fold change relative to sham-operated controls to facilitate robust comparisons across multiple gel membranes.

### 2.8 Immunofluorescence staining and quantitative image analysis

To evaluate the spatial distribution and cellular localization of key mitochondrial and signaling proteins, cryosectioned brain slides were processed for multiplex immunofluorescence staining. Non-specific binding sites were blocked by incubating the tissue sections with 5% normal goat serum (Abcam, Waltham, MA, USA) and 1% bovine serum albumin (BSA; MilliporeSigma, Burlington, MA, USA) in 0.1 M PBS for 1 hour at room temperature. The sections were then incubated overnight at 4°C with primary antibodies directed against CGRP, phosphorylated NRF2 (p-NRF2), microtubule-associated protein 2 (MAP2), PTEN-induced kinase 1 (PINK1), or cytochrome c (**Supplementary Table 2**). Following three washes in PBS, the slides were incubated for 1 hour at room temperature with corresponding fluorophore-conjugated secondary antibodies (**Supplementary Table 2**). For cellular co-localization, sections were sequentially co-stained with either anti-NeuN (to identify neuronal somas) or anti-TOM20 (to label the outer mitochondrial membrane) along with their respective secondary antibodies for 1 hour at room temperature. Nuclei were counterstained with DAPI, and the slides were mounted using Vectashield Antifade Mounting Medium (Vector Laboratories, Burlingame, CA, USA). High-resolution, whole-brain coronal images were captured using an EVOS M7000 imaging system (Thermo Fisher Scientific, Waltham, MA, USA) equipped with a 20× objective and an automated XY-stage stitching function. Quantitative analysis of the immunofluorescence signal was conducted semi-automatically using ImageJ software (NIH, Bethesda, MD, USA). To control for background fluorescence, signal intensity was measured on a standard 8-bit scale (0 to 255) and is presented as reciprocal intensity in arbitrary units (AU), calculated by subtracting the background mean gray value from the target signal mean grey value. Cellular-level expression of CGRP, PINK1, cytochrome c, and p-NRF2 was quantified using a threshold-based method to calculate the positive area of each target marker localized within the somas of 20 randomly selected, NeuN-positive (NeuN^+^) neurons per region of interest (ROI), using three distinct ROIs per anatomical region per animal. Conversely, MAP2 and TOM20 expression profiles were assessed qualitatively to characterize dendritic morphological integrity, mitochondrial structural networks, and spatial interactions with their respective co-expressed targets.

### 2.9 Histopathological assessment of cortical lesion volume

To morphologically evaluate the extent of tissue loss and cortical damage following CCI, the mounted coronal sections were subjected to standard hematoxylin and eosin (H&E) staining. High-resolution, digital whole-slide images of the stained sections were acquired using a PathScan Enabler 5 slide scanner (Meyer Instruments, Houston, TX, USA). Quantitative lesion analysis was performed using ImageJ software (NIH, Bethesda, MD, USA) by an investigator blinded to the experimental cohorts. The boundaries of the necrotic tissue cavity (lesion area), characterized by the loss of cellular architecture and tissue staining, were manually outlined on each section. The total lesion volume for each animal was calculated using the Cavalieri method by integrating the cross-sectional lesion areas across the 28 to 32 serial coronal sections obtained at 400-µm intervals. To control for confounding physiological factors such as post-traumatic brain swelling (edema) or tissue shrinkage during processing, the absolute lesion volume was normalized to the total volume of the contralateral hemisphere and expressed as a percentage value to ensure accurate inter-animal comparisons.

### 2.10 Statistical analysis

All quantitative data are reported as mean ± standard deviation (SD), with individual data points plotted on all graphs to ensure full transparency. Sample sizes were determined by a priori power analysis (α=0.05, β=0.2) based on previous studies, which established a minimum of n=6 per group for biochemical (Western blot) and histological (immunofluorescence, H&E) assays. To account for the higher variability inherent in neurobehavioral testing, the full cohort of n=12 animals per group surviving to 72 hours underwent functional assessments prior to tissue collection. The normality of all datasets was verified using the Shapiro-Wilk and Kolmogorov-Smirnov tests, and homoscedasticity (equality of variances) was assessed using the Brown-Forsythe test. For *Experiment 1*, which evaluated temporal and regional changes across multiple recovery timepoints, data were analyzed using two-way analysis of variance (ANOVA) with treatment condition and survival timepoint as the independent factors. For *Experiment 2* (CGRP receptor blockade studies at 4 and 72 hours), as well as the 72-hour behavioral and lesion volume measurements, statistical differences among the experimental groups were evaluated using ordinary one-way ANOVA. When a significant main effect or interaction was detected by ANOVA, post-hoc pairwise comparisons were performed using Tukey’s honestly significant difference (HSD) multiple comparisons test. In cases where data did not meet normality assumptions, non-parametric equivalents, specifically the Kruskal-Wallis test followed by Dunn’s post-hoc test, were applied. All statistical calculations and graphical representations were generated using GraphPad Prism software (Version 9.0; GraphPad Software, San Diego, CA, USA). For all statistical tests, significance was defined a priori at P < 0.05.

## 3. Results

### 3.1 TNS prevents progressive infarct expansion and functional impairment following TBI

To evaluate the therapeutic efficacy of immediate post-traumatic TNS, which acts as a physical trigger to recruit endogenous oxygen-conserving reflexes [25], we first assessed its impact on progressive tissue loss and neurological deficits. Histological evaluation confirmed that though both vehicle- and TNS-treated groups sustained equivalent primary injuries, as early as 4-hours post-TBI, the cortical infarct volumes began to deviate (9.45±1.28% vs. 7.27±1.03%, respectively; **Fig. 1A**). By 72 hours, the untreated vehicle group showed a substantial progression of secondary tissue damage, with the infarct volume expanding to 13.22±1.46%. Remarkably, acute TNS application arrested this progressive expansion, limiting the 72-hour infarct volume to 8.82±1.15%, which represents a significant 33% reduction in secondary tissue loss.

**Figure 1.**
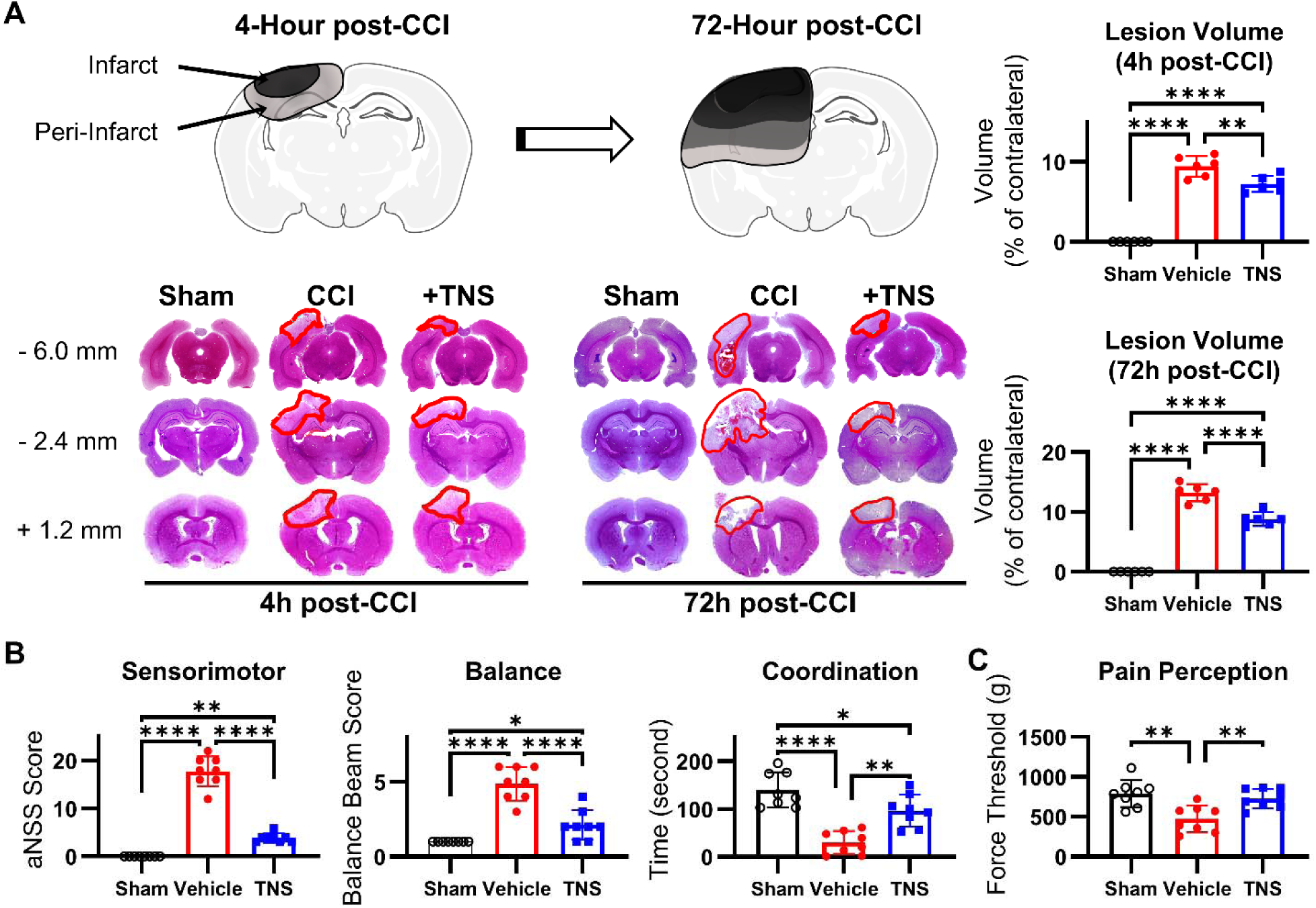
TNS confers structural and functional neuroprotection after TBI. **(A)** Representative H&E stained coronal brain sections at 4h and 72h post-injury, with corresponding quantification of cortical infarct volume. TNS significantly arrested the progressive expansion of the infarct seen in vehicle-treated animals. **(B)** Assessment of sensorimotor function, balance and motor coordination at 72h post-injury. TNS treatment markedly reduced the deficits observed in vehicle-treated animals. **(C)** Evaluation of post-traumatic mechanical pain sensitivity. TBI induced significant pain hypersensitivity in the vehicle-treated group, an effect that was significantly prevented by TNS treatment, maintaining the pain threshold at sham levels.

This structural preservation translated directly into profound neurobehavioral protection. TNS treatment resulted in over 75%, 50%, and 40% reductions in general sensorimotor, balance deficits, and preserving motor coordination (**Fig. 1B**), respectively. Furthermore, while vehicle-treated TBI animals developed severe post-traumatic pain hypersensitivity, demonstrated by a 40% decrease in mechanical pain threshold relative to sham controls, animals receiving TNS exhibited virtually no change in pain sensitivity, remaining comparable to the sham group (**Fig. 1C**). These initial findings demonstrate that early physical neuromodulation via TNS does not merely manage symptoms but actively halts the anatomical and functional progression of secondary brain injury.

### 3.2 TNS mitigates progressive neuronal dysfunction and cytoskeletal decay following TBI

To understand the cellular changes underlying the prevention of progressive tissue loss, we examined neuronal survival (NeuN) and cytoskeletal integrity (MAP2) across a high-resolution timeline (1, 4, 24, and 72 hours) within both the infarct core, the peri-infarct penumbra, and remote contralateral cortex. In vehicle-treated TBI animals, the infarct core showed an immediate, severe reduction in NeuN protein expression starting at 1 hour and worsening continuously through 72 hours (**Fig. 2A**). In the peri-infarct penumbra, this neuronal loss was slightly delayed, with a significant drop appearing at 4 hours and persisting through 24 hours. This decline in survival was accompanied by a progressive loss of the dendritic marker MAP2 in both regions (**Fig. 2A, Supplementary Fig. 1A**). Notably, while MAP2 decreased early in the peri-infarct zone, it exhibited an abnormal, dysregulated increase at 72 hours, indicating severe dendritic breakdown paired with the pathological appearance of MAP2+ reactive astrocytes [47,48].

**Figure 2.**
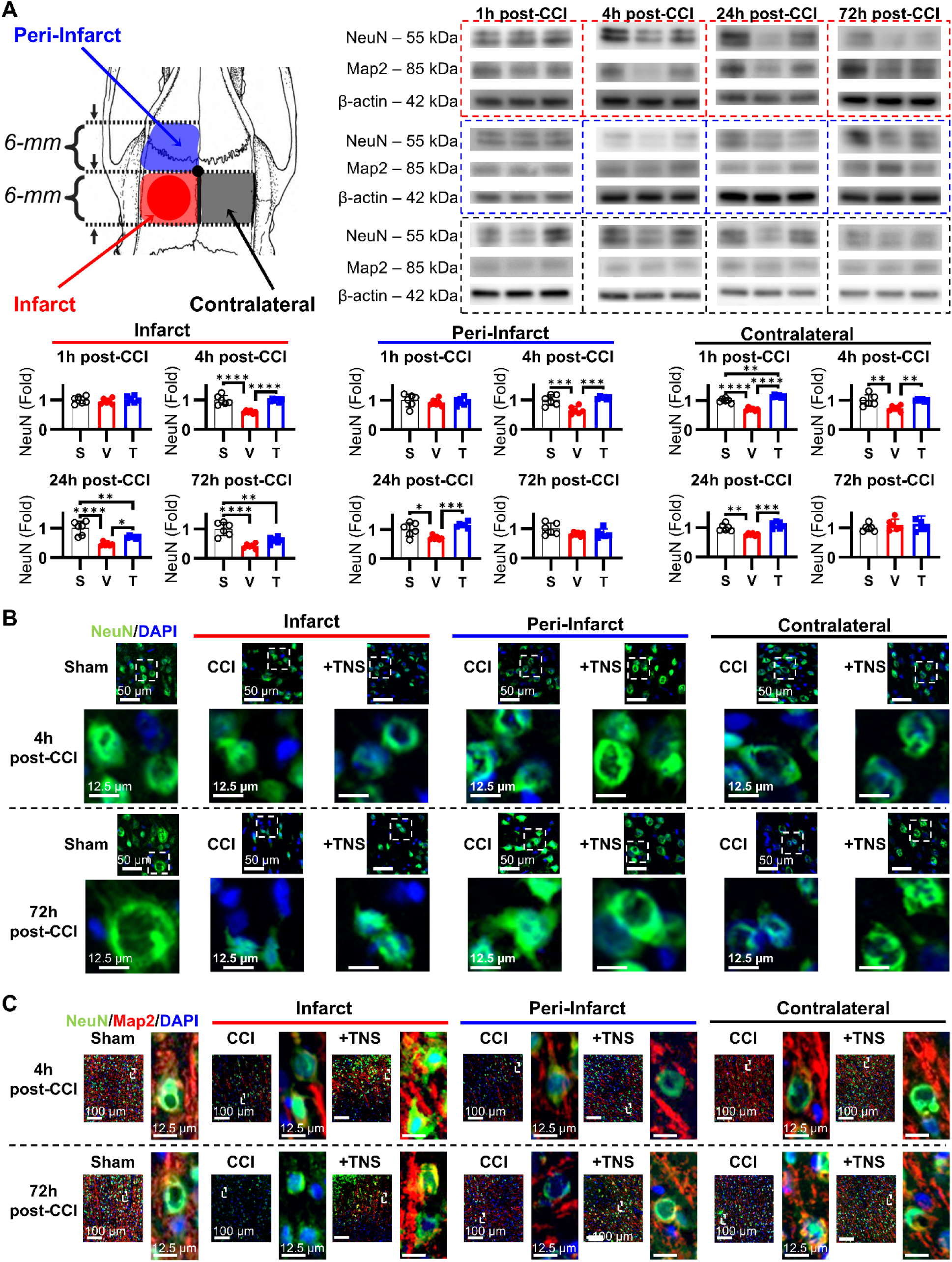
TNS protects neuronal and cytoskeletal structure after TBI. **(A)** Quantification of NeuN (neuronal survival) and MAP2 (dendritic structure) protein expression in the infarct core, peri-infarct penumbra, and contralateral cortex over 72 hours. TNS treatment significantly delayed neuronal loss and preserved cytoskeletal integrity compared to vehicle-treated controls. **(B)** Representative immunofluorescence images of neuronal soma (NeuN, green) at 4h and 72h post-TBI. TNS preserved healthy neuronal morphology and prevented the cellular shrinkage seen in vehicle-treated animals. **(C)** Representative immunofluorescence images of dendritic architecture (MAP2, red) at 4h and 72h post-TBI. TNS maintained intact, organized dendritic processes, particularly in the peri-infarct penumbra, in contrast to the fragmented structures in the vehicle group. (S=sham, V=vehicle, T=TNS-induced diving reflex)

TNS treatment fundamentally shifted these degenerative kinetics, demonstrating a remarkable capacity to protect the structural neuronal matrix. In TNS-treated animals, significant NeuN loss within the infarct core was delayed until 24 hours, while NeuN expression in the peri-infarct penumbra remained largely preserved throughout the entire 72-hour study period. Furthermore, TNS completely prevented the loss of MAP2 in the peri-infarct zone and significantly reduced its depletion within the infarct core, a tissue compartment typically considered unsalvageable.

Immunofluorescence imaging confirmed these molecular changes at the single-cell level. In vehicle-treated animals, neurons in both the core and penumbra showed progressive somatic deformation, shrinking, and fragmentation as early as 1 hour, culminating in near-complete neuronal die-off in the core by 72 hours (**Fig. 2B, Supplementary Fig. 1B**). In contrast, TNS application preserved healthy, rounded neuronal morphology within the peri-infarct region and prevented widespread cell death in the core. Crucially, TNS-treated brains retained straight, largely-intact, and well-organized MAP2+ dendritic processes within the peri-infarct and, to a lesser degree, within the infarct, whereas untreated brains showed severely fragmented, twisted, and degenerated dendritic structures within the peri-infarct and a loss of MAP2 within the infarct (**Fig. 2C**). Significantly, untreated brains exhibited a degree of MAP2 deformation and loss within the contralateral cortex by 72-hours, a pattern which was wholly missing with TNS treatment. These results demonstrate that early physical neuromodulation preserves not only individual cell survival but also the structural architecture of the neuronal networks required for functional recovery.

### 3.3 TNS drives an early, sustained neuronal CGRP surge that correlates with regional cell survival

To determine whether CGRP is a primary mediator of the observed neuronal preservation, we examined both overall tissue CGRP and neuron-specific CGRP levels across the three cortical regions over the 72-hour timeline. Western blot analysis of the infarct core revealed that TBI triggered an initial increase in overall tissue CGRP at 1-hour post-injury. TNS application significantly amplified this response, driving a more than 2-fold increase in CGRP relative to the vehicle group by 4 hours, which rose to a more than 4-fold increase by 24 hours (**Fig. 3A**). In the peri-infarct penumbra, the natural CGRP response in vehicle-treated animals was critically delayed, only rising at 24 hours. TNS completely eliminated this delay, triggering a rapid and significant CGRP increase as early as 4 hours. In the contralateral cortex, vehicle-treated animals showed a temporary drop in CGRP at 4 hours, which was prevented by TNS. Both groups eventually showed an increase in contralateral CGRP at 72 hours, likely reflecting a delayed, brain-wide response to secondary injury.

**Figure 3.**
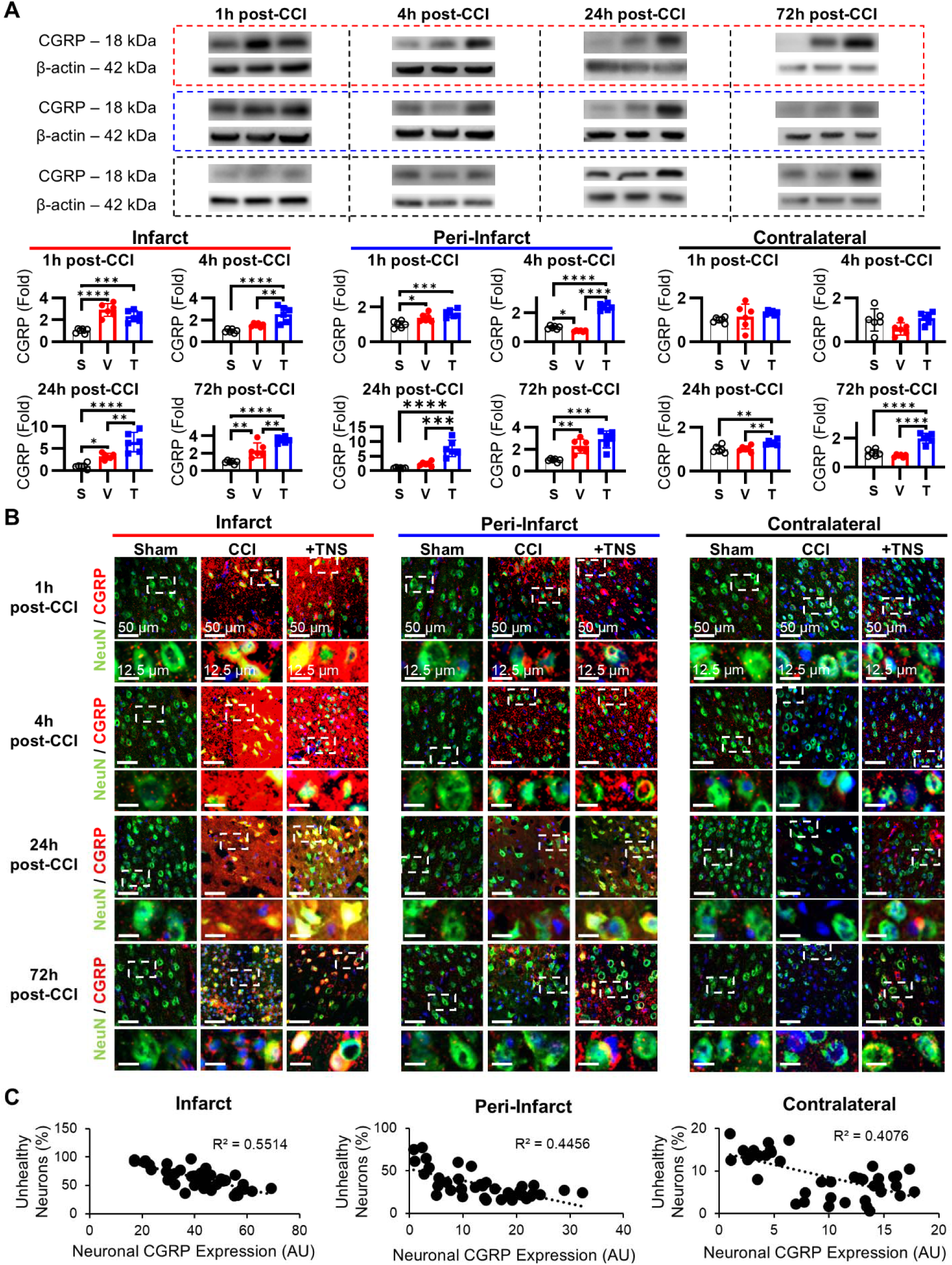
**TNS accelerates the CGRP response, which is essential for neuronal survival**. **(A)** Quantification of total tissue CGRP levels across cortical regions. TNS induced a more rapid and robust CGRP increase in the infarct core and peri-infarct penumbra compared to the delayed endogenous response in vehicle-treated animals. **(B)** Representative immunofluorescence images and quantification of neuronal CGRP. TNS treatment led to a sustained elevation of CGRP within neurons, preventing the depletion observed in the vehicle group. **(C)** Correlation analysis reveals a strong positive relationship between regional CGRP levels and neuronal survival (NeuN), demonstrating that higher CGRP levels are associated with greater cell survival. (S=sham, V=vehicle, T=TNS-induced diving reflex)

Importantly, our analysis revealed a clear difference between overall tissue CGRP and intra-neuron CGRP (**Fig. 3B, Supplementary Fig. 2**). In the infarct core of untreated vehicle animals, neuronal CGRP showed only a mild, transient increase before falling significantly by 72 hours, despite the nigh-on immediate increase in tissue CGRP expression. In contrast, TNS sustained a progressive increase in neuronal CGRP throughout the first 24 hours, keeping it highly elevated through 72 hours. Furthermore, while contralateral neuronal CGRP was unaffected in vehicle animals, TNS-treated animals showed a marked, brain-wide increase in neuronal CGRP. This broad upregulation is highly significant because, despite CGRP’s classical association with pain transmission [30], these TNS-treated animals did not develop pain hypersensitivity, pointing to a non-pain-inducing, protective role for neuromodulated CGRP in injurious contexts.

These temporal dynamics highlight a critical therapeutic window: the brain’s natural, late-stage CGRP increase at 24 to 72 hours in untreated animals occurs long after neurons have begun to deform and die. By forcing an immediate and sustained intracellular CGRP surge, TNS acts within this critical early window to protect vulnerable cells. This direct relationship is confirmed by correlation analysis between CGRP levels and neuronal health (**Fig. 3C**). We found a strong, region-specific correlation showing that higher CGRP levels directly predict greater neuronal survival, with the strongest correlation occurring in the most severely threatened areas (Infarct core: R^2^ = 0.5514; Peri-infarct penumbra: R^2^ = 0.4456; Contralateral cortex: R^2^ = 0.4076). This strong link suggests that early CGRP availability is a central driver of post-traumatic neuronal survival.

### 3.4 TNS stabilizes the kinetics and neuronal expression of the PINK1/Parkin pathway following TBI

To determine whether TNS-mediated neuroprotection is driven by the preservation of mitochondrial quality control, we analyzed key markers of the mitophagy and autophagy pathways, PINK1, Parkin, and LC3II, using Western blot and immunofluorescence imaging across our established 72-hour timeline.

Within the infarct core of untreated TBI animals, Western blot analysis revealed an immediate, severe reduction in the mitochondrial damage-sensor PINK1 starting at 1-hour post-injury and persisting through 72 hours (**Fig. 4A**). This decline occurred alongside a rapid, reactive spike in Parkin expression. Because LC3II levels remained largely unchanged, this mismatched pattern, characterized by upstream PINK1 depletion coupled with paradoxical downstream Parkin upregulation, suggests a severe post-traumatic uncoupling or dysregulation of the mitochondrial quality-control machinery. TNS intervention successfully rescued this signaling pathway, preventing the loss of PINK1 within the infarct core and further potentiating Parkin expression starting at 4 hours post-TBI. In the peri-infarct penumbra of vehicle-treated animals, the endogenous mitochondrial response was critically delayed: PINK1 exhibited a transient decrease followed by a late, reactive increase at 24 hours, while Parkin expression did not rise until 24 hours. TNS largely eliminated this kinetic delay, preventing the early decline in PINK1 and accelerating its protective upregulation to 4 hours post-injury. This temporal advancement was accompanied by a rapid, sustained increase in Parkin starting as early as 1 hour, demonstrating that TNS rapidly mobilizes and stabilizes the PINK1/Parkin pathway during the critical hyper-acute window of secondary injury.

**Figure 4.**
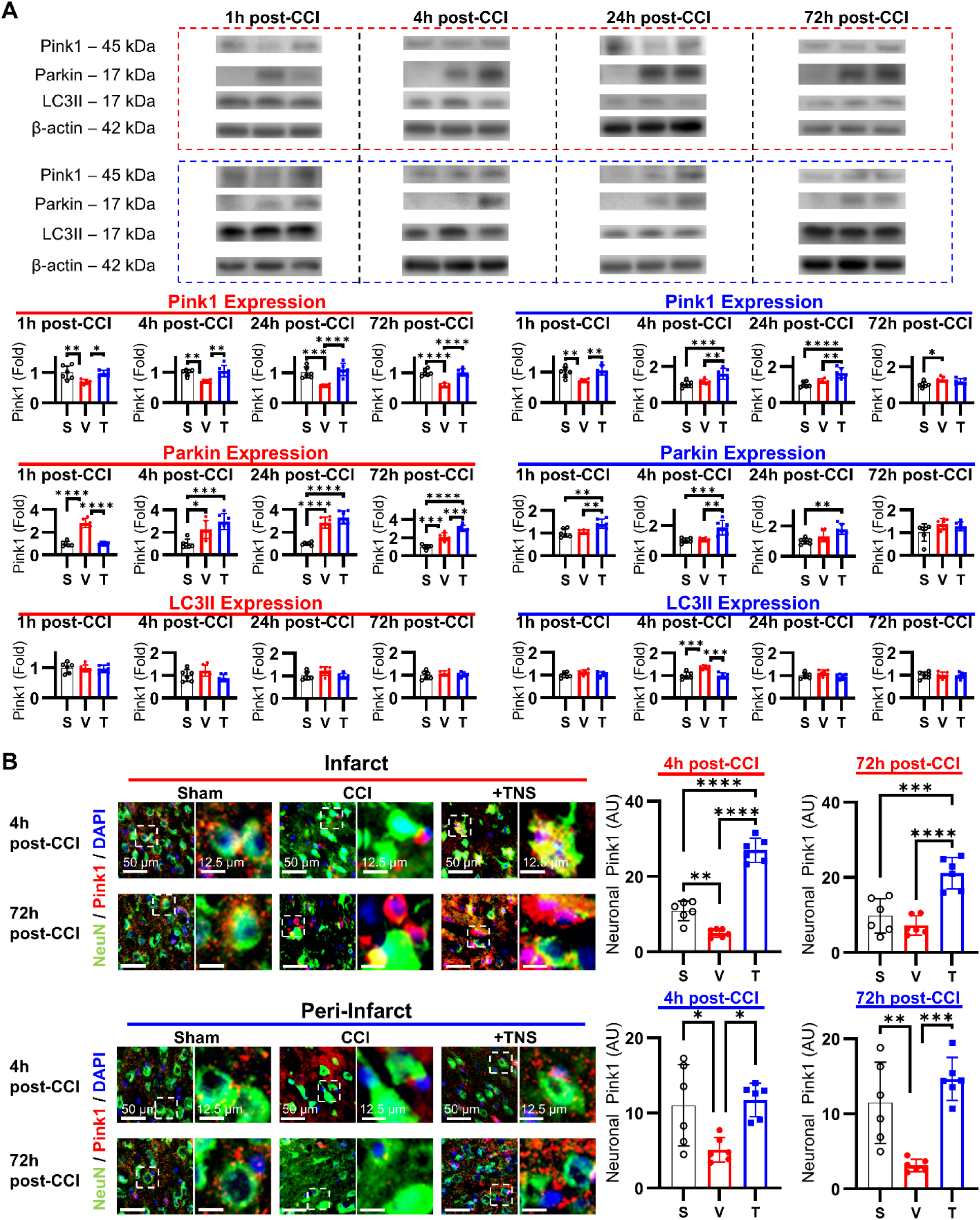
TNS rescues post-traumatic dysregulation of the PINK1/Parkin pathway. **(A)** Western blot quantification of key mitophagy proteins at 1h, 4h, 24h, and 72h post-injury. TNS prevented the acute loss of the mitochondrial damage-sensor PINK1 and accelerated the upregulation of both PINK1 and Parkin in injured cortical regions, restoring coordinated pathway activation compared to the dysregulated response in vehicle-treated animals. **(B)** Immunofluorescence imaging of neuronal PINK1 (red) and NeuN (green) at 4h and 72 post-injury. TNS preserved the intracellular localization of PINK1 within neurons and prevented the pathological aggregation that was prominent in vehicle-treated animals at 72 hours post-injury. (S=sham, V=vehicle, T=TNS-induced diving reflex)

To complement these biochemical findings, we performed immunofluorescence imaging at 4 hours (hyper-acute) and 72 hours (sub-chronic) post-injury to evaluate spatial and cellular expression patterns (**Fig. 4B**). In vehicle-treated TBI animals, neuronal PINK1 expression within the infarct core was severely reduced at 4 hours. By 72 hours, although a nominal return of PINK1 signal was observed, it appeared highly aggregated and failed to colocalize with NeuN-positive neurons. This pathological, extracellular aggregation suggests a loss of functional, cell-autonomous mitochondrial quality control. A similar, though less pronounced, depletion of neuronal PINK1 was observed in the untreated peri-infarct penumbra. In contrast, TNS treatment preserved robust, intracellular neuronal PINK1 expression in both the core and penumbra at both 4 and 72 hours. Notably, TNS-treated brains showed no signs of the pathological extracellular PINK1 clumping observed in the vehicle group, displaying instead a diffuse, structured distribution pattern resembling that of sham controls.

These findings indicate that TBI disrupts the coordinated activation of mitochondrial quality control, creating an uncoupled state where reactive Parkin upregulation cannot be functionally utilized due to the rapid depletion and succeeding pathological agglomeration of its upstream sensor, PINK1. By preventing the hyper-acute loss of neuronal PINK1 and accelerating downstream Parkin kinetics, TNS resolves this signaling mismatch. These results provide initial indications that TNS restores mitophagic homeostasis, shifting the post-traumatic microenvironment from a state of failed organelle clearance toward coordinated, active mitochondrial preservation during the critical early window of secondary neurodegeneration.

### 3.5 TNS resolves a post-traumatic TOM20 structural bottleneck to facilitate PINK1 mitochondrial docking

In order to further examine the mechanism underlying the observed destabilization of the PINK1/Parkin pathway in TBI, characterized by the late-stage increases and agglomeration, and the stabilization thereof by TNS, we examined the organization of the outer mitochondrial membrane import receptor, TOM20, and its spatial relationship with PINK1. Under physiological stress, functional initiation of the mitophagy cascade requires PINK1 to dock with TOM20 to assemble the PINK1-TOM-TIM23 import complex [42,49–52]. However, this signaling axis requires precise stoichiometric and structural organization of TOM20. Pathological aggregation of this receptor can disrupt outer membrane import geometry, physically preventing PINK1 from stable membrane docking and potentially explaining the observed extra-neuronal agglomeration of PINK1 and the ostensibly ineffectual late-stage increase of Parkin.

In sham-injured brains, immunofluorescence imaging revealed that TOM20 was distributed in a highly organized, diffuse, and punctate pattern throughout the cytoplasm, reflecting a healthy, homeostatic mitochondrial network (**Fig. 5, Supplementary Fig. 3**). Following TBI, vehicle-treated brains exhibited severe, pathological TOM20 aggregation and clumping within both the infarct core and the peri-infarct penumbra. High-resolution spatial analysis showed that these abnormal TOM20 aggregates were physically segregated from the remaining PINK1 signal in vehicle-treated animals. Microdomains characterized by heavy, clumped TOM20 were nearly devoid of PINK1 expression. This spatial mismatch suggests a post-traumatic cessation of PINK1 mitochondrial import, possibly induced by pathological TOM20 accumulation and mitochondrial membrane dysfunction preceding (1) loss of functional interaction between the PINK1 N-terminal–C-terminal extension module and the cytosolic domain of the Tom20 subunit of the TOM complex and (2) the impairment or loss of TIM23, both of which disenable the bound PINK1 from functionally importing into the mitochondria [42,49–52]. Accumulated PINK1 within the cytosol cannot bind to Parkin and is then degraded by either the ubiquitin-proteasome system or aggresome formation [53], leading to the observed mismatch in PINK1 and TOM20 locality. TNS intervention restricted the pathological clumping of TOM20 to a transient appearance within both the infarct core and peri-infarct which peaked at 4 hours post-injury which then proceeded to decrease, apparently having prevented the persistence of the possible structural bottleneck. By 72 hours, TOM20 distribution in both the infarct and peri-infarct regions returned to a diffuse, punctate pattern resembling that of sham controls, which coincided with the re-establishment of spatial overlap between TOM20 and PINK1. Of chief import though, was the observance of co-localization of PINK1 with the mildly aggregated TOM20 in TNS, indicating a likely intact functioning of the PINK1-TOM20 binding and mitochondrial processing.

**Figure 5.**
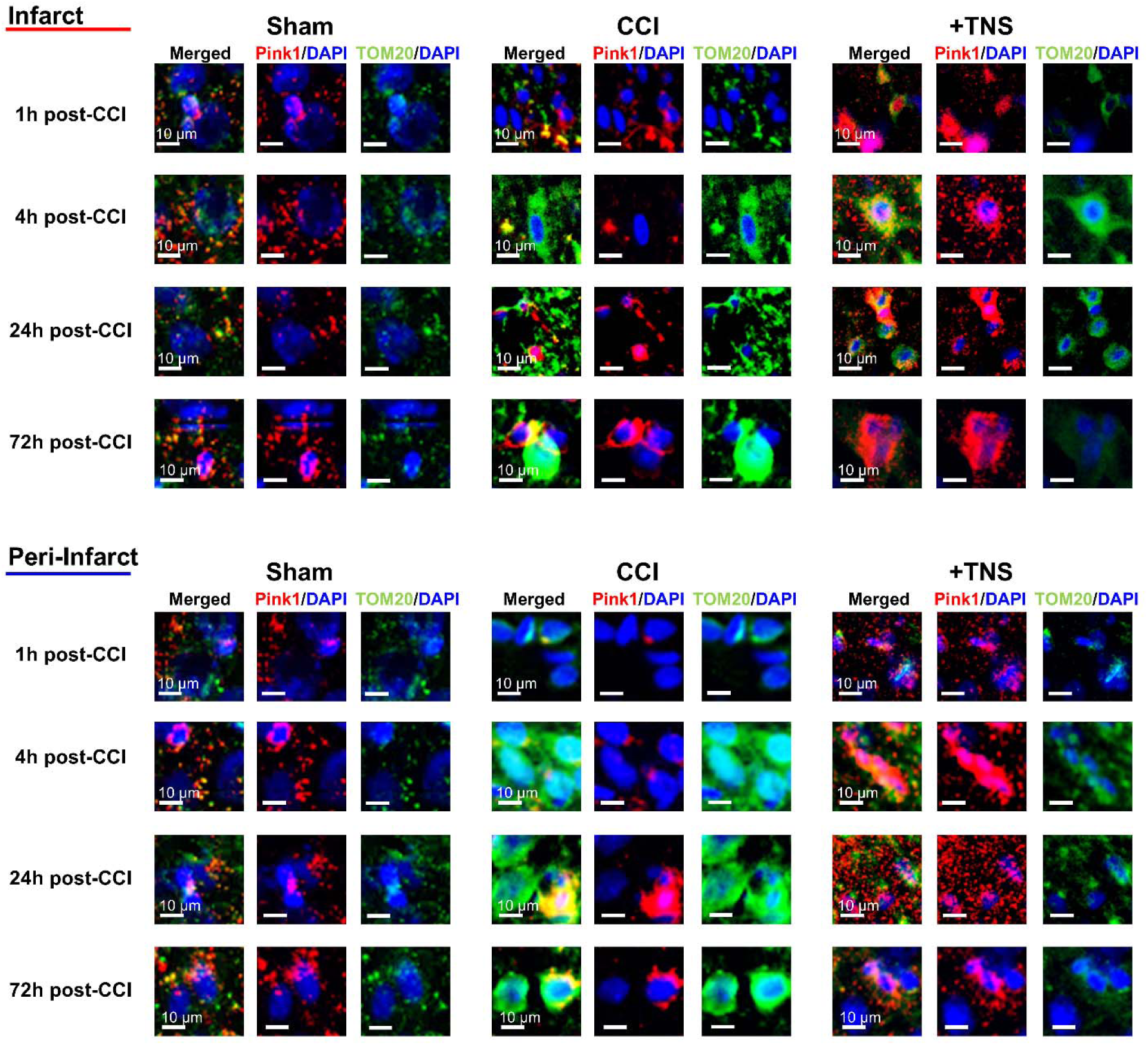
TNS preserves outer mitochondrial membrane architecture and PINK1 co-localization. Representative immunofluorescence images of the mitochondrial import receptor TOM20 (green) and PINK1 (red) at 1h, 4h, 24h, and 72h post-injury. In vehicle-treated animals, TBI induces pathological aggregation of TOM20 and its spatial segregation from PINK1, physically obstructing the quality control pathway. TNS treatment prevents this persistent aggregation, maintaining a diffuse, physiological distribution of TOM20 and ensuring its co-localization with PINK1.

These findings provide preliminary indications suggesting a novel structural mechanism of post-traumatic mitochondrial pathology characterized by the physical aggregation of TOM20, which acts as a molecular bottleneck to block PINK1 docking and arrest mitochondrial rescue. By preserving the diffuse, physiological geometry of TOM20 on the outer mitochondrial membrane, TNS preserves the physical docking interface required for PINK1 recruitment. This likely structural salvage indicates a potentially key mechanism by which physical neuromodulation maintains mitochondrial signaling integrity and protects vulnerable neuronal networks from progressive metabolic collapse.

### 3.6 CGRP inhibition triggers pathological TOM20 aggregation and uncouples the PINK1/Parkin axis

To determine whether the observed modulation of mitochondrial quality control is related to CGRP signaling, we evaluated the effects of pharmacological CGRP receptor inhibition in the context of TBI.

Western blot analysis revealed that pharmacological inhibition of CGRP signaling during TBI led to a significant decrease in PINK1 expression within the peri-infarct penumbra (**Fig. 6A**). Interestingly, this decrease in PINK1 was accompanied by an increase in total Parkin expression, echoing and pre-empting the dysregulation observed in the vehicle animals. Because Parkin requires PINK1-mediated phosphorylation for translocation and functional activation on the outer mitochondrial membrane [54,55], this mismatched pattern, characterized by depleted PINK1 alongside elevated Parkin, indicates a further exacerbation and uncoupling of the pathway leading to a non-functional reactive response.

**Figure 6.**
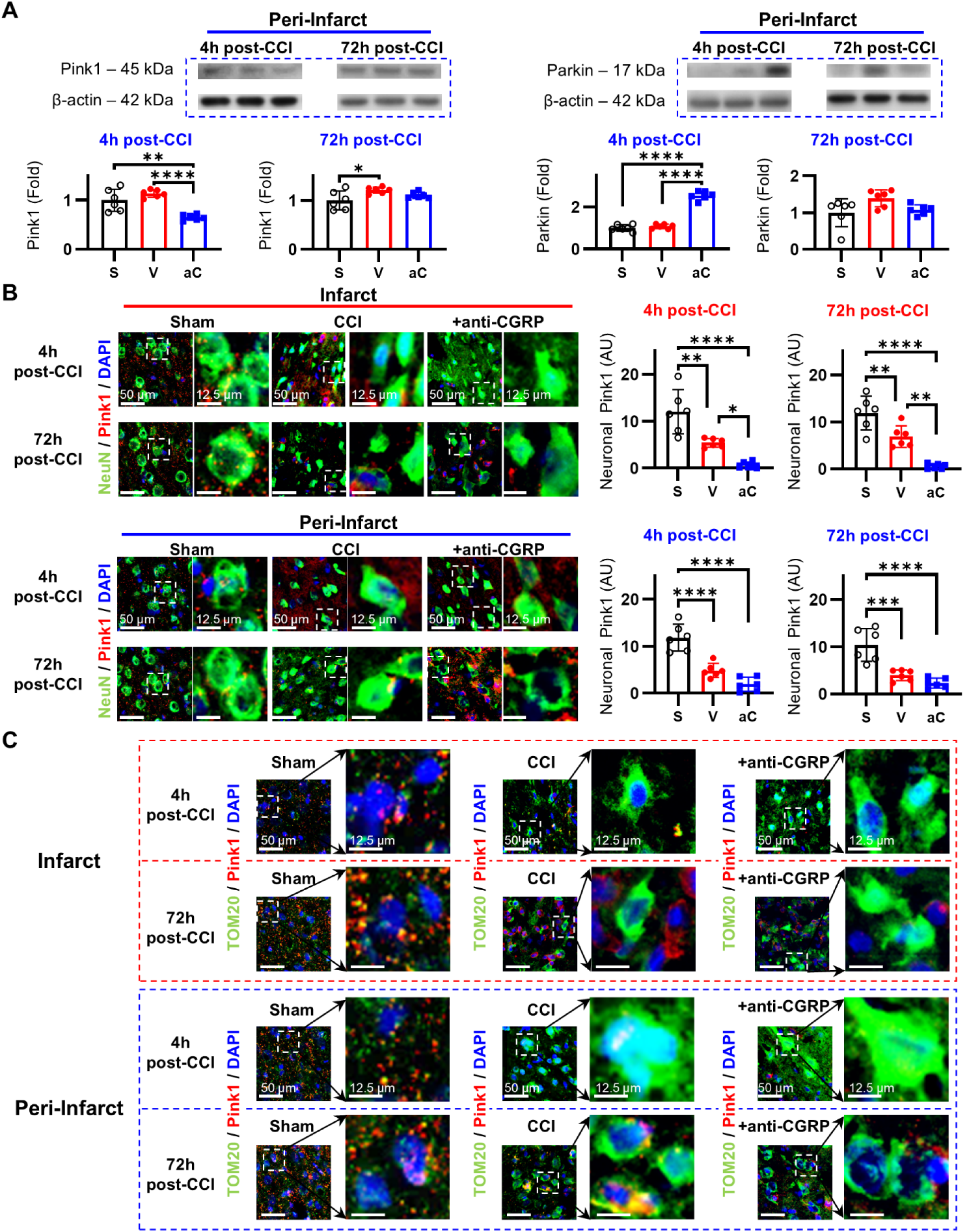
CGRP signaling is required to maintain mitochondrial structural and signaling integrity. **(A)** Quantification of cortical PINK1 and Parkin expression following CGRP receptor inhibition at 4h and 72h post-injury. CGRP blockade induced an uncoupled response, characterized by depleted PINK1 in conjunction with elevated Parkin levels, indicating pathway failure. **(B)** Immunofluorescence images showing that CGRP inhibition exacerbates the loss of PINK1 (green) from neurons after TBI. **(C)** CGRP blockade resulted in severe aggregation of the mitochondrial protein TOM20 (green), preventing its co-localization with PINK1 (red) and structurally disrupting the mitochondrial rescue pathway. (S=sham, V=vehicle, aC=anti-CGRP)

Immunofluorescence analysis provided a striking structural explanation for this signaling failure. CGRP receptor inhibition was shown to exacerbate the post-traumatic loss of neuronal PINK1 within both the infarct core and the peri-infarct penumbra (**Fig. 6B**). Crucially, this was accompanied by a severe derangement of gating mechanisms along the outer mitochondrial membrane. TOM20 staining in CGRP-inhibited animals revealed highly accentuated, persistent aggregation of the receptor (**Fig. 6C**). This structural disruption is likely to have underwritten the complete prevention of spatial colocalization between TOM20 and the remaining PINK1, suggesting that CGRP signaling is necessary to maintain the physical interface required for mitophagy initiation.

These loss-of-function experiments establish CGRP as a likely crucial upstream regulator of the mitochondrial quality control machinery after TBI. Blocking CGRP signaling triggers a catastrophic failure at the outer mitochondrial membrane, precipitating the pathological aggregation of TOM20 and preventing the physical docking of PINK1. This intimates that endogenous CGRP is required to preserve the structural integrity and geometry of the outer membrane docking apparatus, thereby providing preliminary identification of the trigeminal-CGRP axis as a necessary biological link between physical neuromodulation and the functional initiation of the PINK1/Parkin pathway.

### 3.7 TNS preserves mitochondrial antioxidant defenses and prevents cytochrome c leakage

To evaluate the downstream functional consequences of this mitochondrial structural salvage, we examined the mitochondrial antioxidant enzyme Superoxide Dismutase 2 (SOD2) and the leakage of cytochrome c from the intermembrane space into the cytoplasm.

Following TBI, untreated vehicle animals exhibited a significant loss of SOD2 expression within both the infarct core and the peri-infarct penumbra (**Fig. 7A**). In the infarct core, SOD2 levels decreased by approximately 30% at 4 hours post-injury and remained suppressed through 72 hours. In the peri-infarct penumbra, SOD2 levels showed a 10% reduction at 4 hours. Acute TNS application successfully prevented this decline, maintaining SOD2 expression levels near sham baselines in both regions over the 72-hour period.

**Figure 7.**
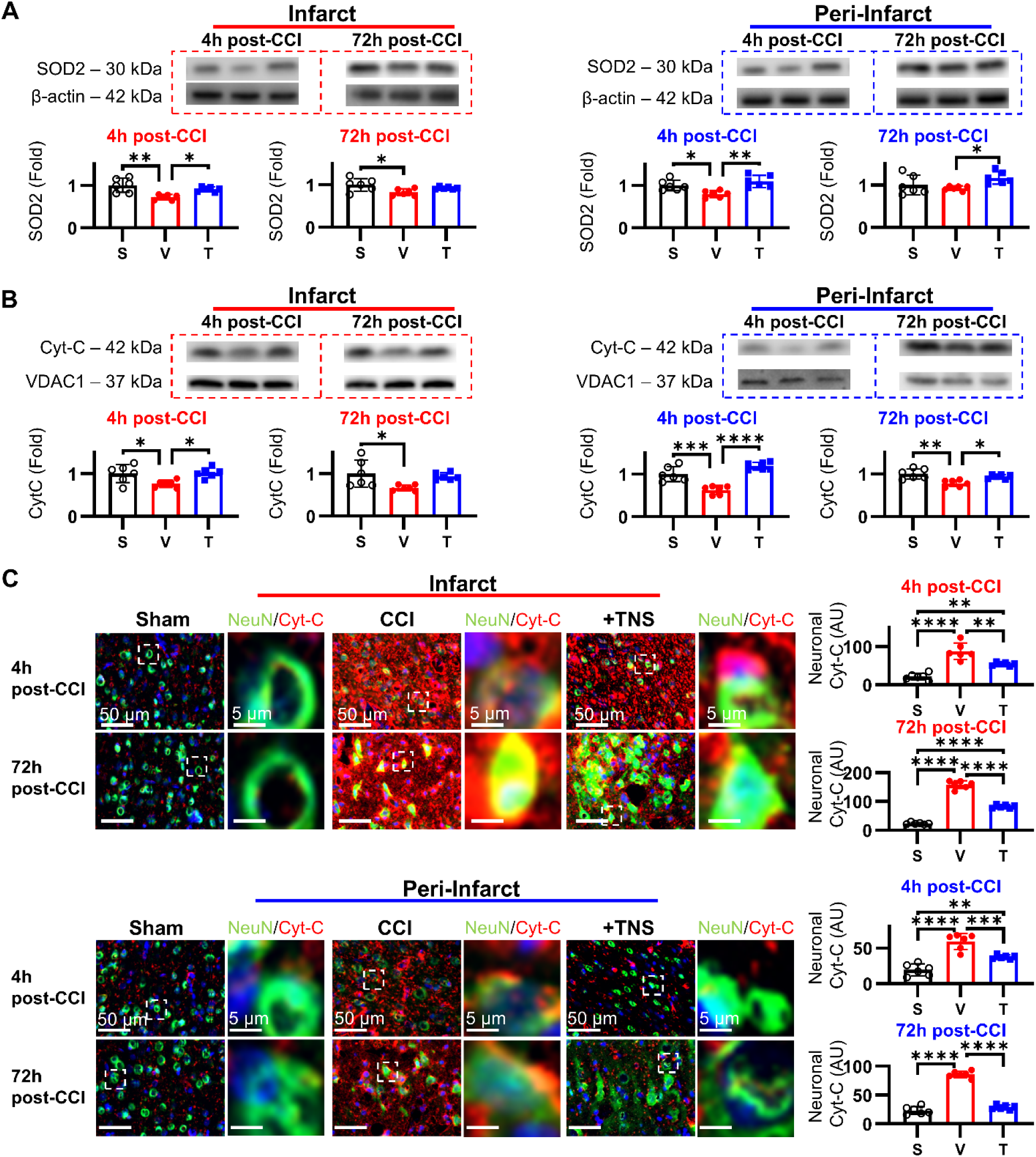
TNS preserves mitochondrial function and prevents pro-apoptotic factor release. **(A)** Quantification of the mitochondrial antioxidant enzyme SOD2. TNS prevented the post-traumatic decline in SOD2 expression observed in vehicle-treated animals. **(B)** Western blot of mitochondrial fractions shows that TNS maintained cytochrome c levels within mitochondria, preventing its TBI-induced depletion. **(C)** Immunofluorescence imaging of cytochrome c. TNS significantly blunted the leakage of cytochrome c into the cytoplasm. This protective effect was dependent on CGRP signaling, as shown by experiments with CGRP inhibition (see also Supplementary Fig. 4). (S=sham, V=vehicle, T=TNS-induced diving reflex)

To assess outer mitochondrial membrane permeability, we monitored the distribution of cytochrome c [56–58]. In vehicle-treated TBI animals, Western blot analysis of isolated mitochondrial fractions revealed a substantial, global decrease in mitochondrial cytochrome c at both 4 and 72 hours post-injury, indicating a breakdown of outer membrane retention capacity (**Fig. 7B**). TNS treatment prevented this loss, sustaining mitochondrial cytochrome c levels and indicating the preservation of outer membrane structural integrity.

Immunofluorescence imaging of brain sections directly demonstrated the CGRP-dependent nature of this outer membrane preservation (**Fig. 7C, Supplementary Fig. 4**). In vehicle-treated TBI animals, neuronal cytoplasmic cytochrome c (representing leaked protein) increased 4-fold within the infarct core at 4 hours post-injury. In the peri-infarct penumbra, CGRP inhibition further exacerbated cytochrome c release, resulting in a sustained 5-fold elevation of cytoplasmic cytochrome c at 72 hours, indicating progressive, unmitigated mitochondrial breakdown. In contrast, TNS application significantly blunted this translocation. In TNS-treated animals, the accumulation of cytoplasmic cytochrome c in both the infarct and peri-infarct regions was reduced by more than half compared to the vehicle group, demonstrating that TNS effectively limits the release of pro-apoptotic factors into the neuronal cytoplasm.

These results demonstrate that the structural salvage of mitochondria by TNS translates directly into functional neuroprotection. By sustaining SOD2-mediated antioxidant defenses, TNS preserves outer mitochondrial membrane integrity. This stabilization prevents the catastrophic cytoplasmic leakage of the pro-apoptotic factor cytochrome c, thereby directly linking TNS-mediated neuromodulation to the inhibition of downstream apoptotic pathways.

### 3.8 TNS activates downstream NRF2 antioxidant defenses and mitigates post-traumatic oxidative stress

To determine whether the structural preservation of mitochondria and retention of cytochrome c translate to enhanced cellular defense networks, we examined the activation of NRF2, a master transcriptional regulator of the endogenous antioxidant response, via compartment-specific Western blot and immunofluorescence analyses.

Subcellular fractionation and compartment analysis of the infarct core revealed that TNS treatment stimulated a robust 2.3-fold increase in nuclear NRF2 translocation at the hyper-acute phase, far exceeding the modest 1.3-fold reactive increase observed in vehicle-treated TBI animals (**Fig. 8A**). Within the salvageable peri-infarct penumbra, where untreated TBI animals exhibited a progressive loss of nuclear NRF2, TNS treatment successfully preserved NRF2 activation, preventing more than 50% of the injury-induced depletion.

**Figure 8.**
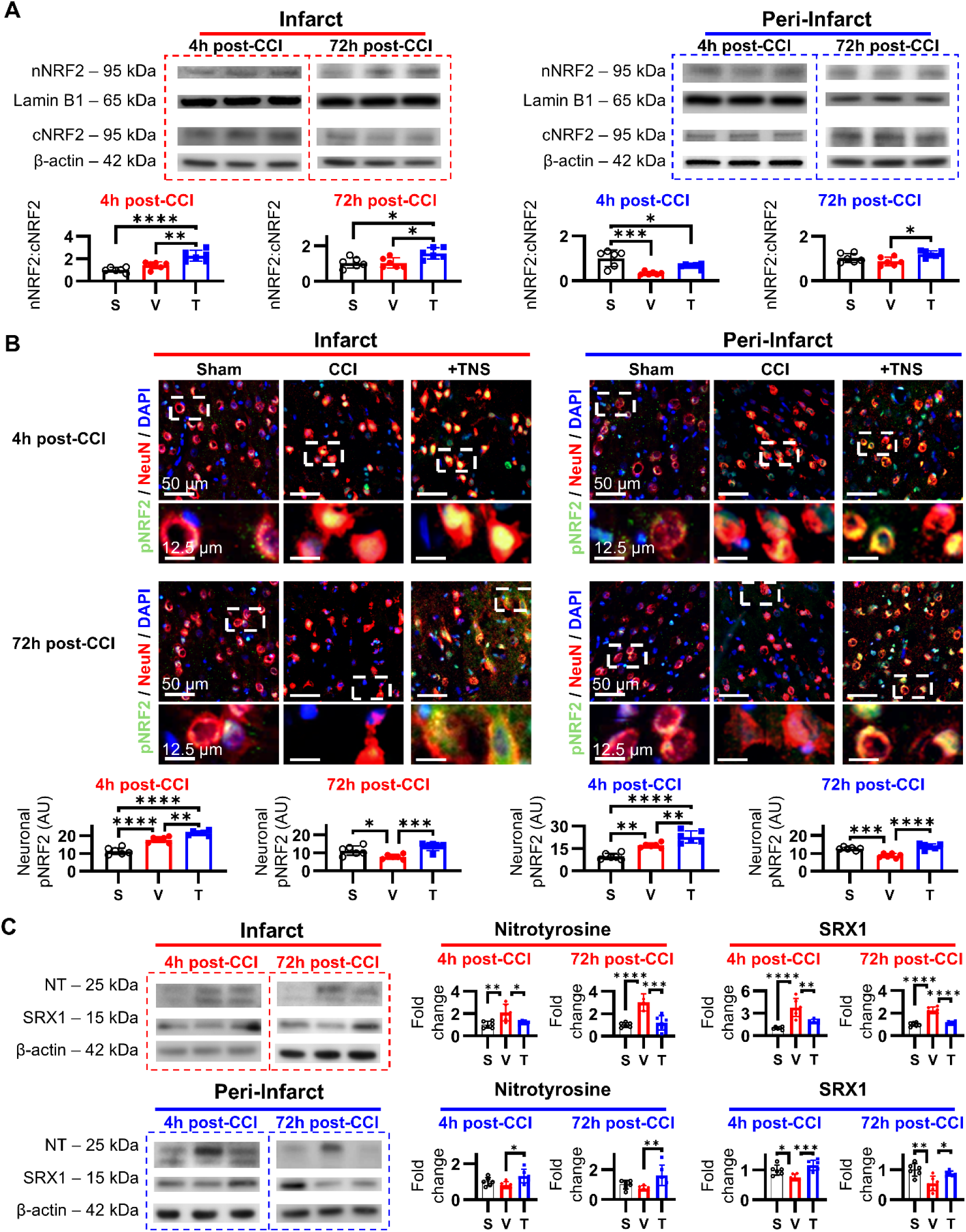
TNS enhances the NRF2 antioxidant response and reduces oxidative stress. **(A)** Western blot analysis of subcellular fractions shows that TNS promotes the nuclear translocation of the master antioxidant regulator NRF2 in injured cortical regions. **(B)** Immunofluorescence for phosphorylated NRF2 (p-NRF2) demonstrates that TNS sustains the activation and nuclear localization of this key transcription factor within neurons. **(C)** TNS treatment mitigated post-traumatic oxidative stress, as shown by reduced nitrotyrosine accumulation and preserved expression of the antioxidant enzyme SRX1 in the peri-infarct penumbra. (S=sham, V=vehicle, T=TNS-induced diving reflex)

To confirm the transcriptional activation of NRF2, we performed immunofluorescence imaging evaluating its active, phosphorylated form (p-NRF2) at 4 and 72 hours post-injury (**Fig. 8B**). In vehicle-treated TBI animals, p-NRF2 expression showed a transient, hyper-acute spike at 4 hours before undergoing a severe depletion by 72 hours, reflecting a failure to sustain antioxidant signaling. In contrast, TNS treatment significantly enhanced p-NRF2 expression at 4 hours and completely prevented its sub-chronic depletion at 72 hours, maintaining robust, localized nuclear expression within surviving neurons across both the core and penumbra.

To evaluate the functional consequence of this sustained NRF2 activation, we measured tissue levels of nitrotyrosine, a classical marker of peroxynitrite-mediated protein nitrosylation and nitrosative stress, and Sulfiredoxin-1 (SRX1), an essential downstream antioxidant enzyme (**Fig. 8C**). In vehicle-treated TBI animals, the peri-infarct penumbra exhibited a hyper-acute and persistent increase in nitrotyrosine accumulation, which was mirrored by a significant depletion of SRX1. TNS treatment successfully reversed these pathological shifts, preventing the accumulation of nitrotyrosine and maintaining robust SRX1 expression within both the infarct and the peri-infarct tissue.

In summary, these final results demonstrate that the mitochondrial rescue mediated by TNS culminates in the activation of a robust, genome-wide cellular defense program. By sustaining NRF2 nuclear translocation and phosphorylation, TNS preserves downstream antioxidant machinery like SRX1. This transcriptional activation effectively quenches the hyper-acute post-traumatic oxidative burst, demonstrating how TNS initiates a significant trans-cellular cascade, stretching from systemic reflexes down to organelle-level structural salvage and genomic antioxidant defenses, to halt progressive neurodegeneration.

## 4. Discussion

This study identifies a direct mechanistic link between non-invasive neuromodulation and the preservation of mitochondrial quality control after TBI. While TNS is known to improve neurological outcomes, the cellular mechanisms linking trigeminal input to organellar preservation have remained unclear. Here, we demonstrate that acute, tuned TNS modulates the CGRP-mediated mitochondrial PINK1/Parkin quality control system. Our findings intimate that TNS-induced CGRP release prevents the pathological aggregation of the outer mitochondrial membrane protein TOM20, a structural failure that otherwise obstructs PINK1 signaling (**Fig. 9**). This preservation of mitochondrial architecture prevents the release of pro-apoptotic cytochrome c and supports NRF2-mediated antioxidant defenses. By identifying CGRP as a potential molecular mediator between TNS and mitochondrial viability, these findings provide a translatable, mechanistically grounded strategy for protecting the bioenergetically compromised penumbra during the critical acute window of brain injury.

**Figure 9.**
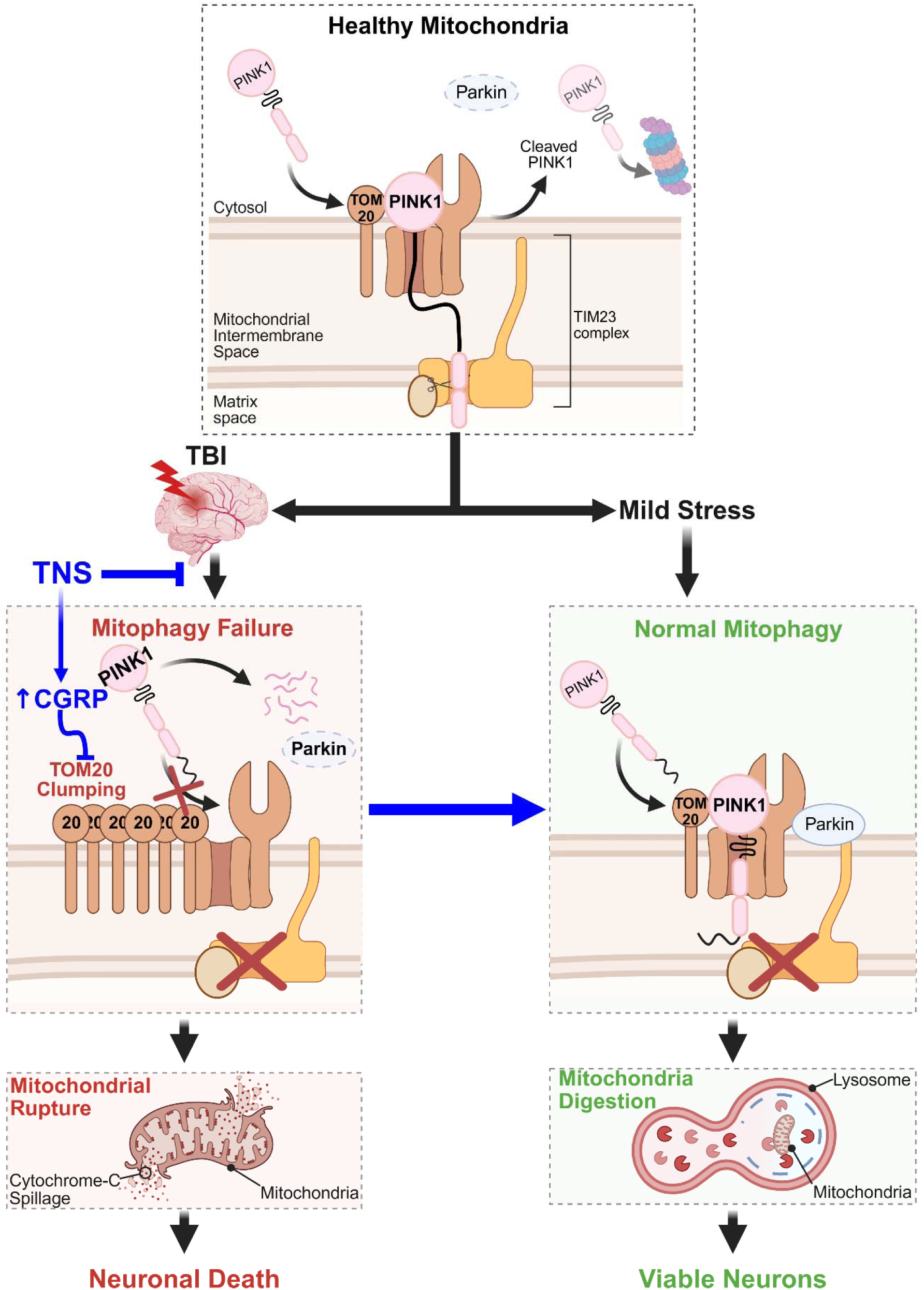
Conceptual framework for TNS-Mediated neuroprotection via mitochondrial structural salvage. This schematic illustrates the central mechanism identified in this study. Following TBI, the endogenous stress response is insufficient to prevent the pathological aggregation of the outer mitochondrial membrane protein TOM20. This structural failure creates a physical bottleneck, obstructing PINK1 docking and arresting the PINK1/Parkin mitochondrial quality control pathway. The resulting mitochondrial dysfunction leads to the release of pro- apoptotic cytochrome c, failure of NRF2-mediated antioxidant defenses, and progressive neurodegeneration. Acute TNS intervention provides a timely and robust CGRP surge. This signal preserves the physiological, diffuse architecture of TOM20, thereby maintaining the physical interface for PINK1 signaling. By keeping the PINK1/Parkin pathway functional, TNS stabilizes mitochondria, prevents cytochrome c leakage, activates NRF2 antioxidant responses, and ultimately halts secondary injury in the vulnerable penumbra.

The spatial and temporal patterns of PINK1 expression after TBI provide critical insights into the progression of secondary injury. Within the injury core, PINK1 levels decreased as early as 1 hour and remained suppressed through 72 hours, reflecting a rapid and severe failure of mitochondrial quality control machinery. In contrast, within the peri-injury penumbra, PINK1 levels showed a delayed increase. This regional heterogeneity demonstrates that the post-traumatic mitochondrial response is not uniform and depends on the local severity and timing of the injury [59,60]. This distinction is therapeutically significant: while the injury core suffers an early, catastrophic failure of mitochondrial signaling, the surrounding penumbra retains the capacity to activate this defense but does so with a critical delay. This peri-infarct penumbra represents a highly salvageable tissue compartment where early intervention can prevent the transition from reversible cellular stress to permanent injury [21,61]. Our data show that TNS effectively modulates both components of this regional response. In the injury core, TNS stabilized PINK1 expression, preventing the persistent loss observed in untreated animals. In the peri-infarct penumbra, TNS accelerated the PINK1 response, advancing the time to significant elevation from 24 hours to 4 hours post-injury. This 20-hour advancement is highly relevant, as mitochondrial decay and cell-death signaling develop rapidly during this early period. Promoting PINK1 function earlier allows the cell to manage damaged mitochondria before the injury cascade becomes irreversible [7,62,63]. Furthermore, TNS protected the contralateral cortex from remote secondary injury, preserving neuronal integrity and maintaining PINK1 signaling. This indicates that focal TBI induces widespread cellular stress, and that TNS activates a broad neuromodulatory network capable of protecting tissue far beyond the primary injury site.

A novel finding of this study is the preliminary identification of a potential structural failure at the outer mitochondrial membrane that physically obstructs the PINK1/Parkin quality control pathway after TBI. We demonstrate that TBI induces the pathological agglomeration of the mitochondrial import receptor, TOM20 [64,65]. This discovery potentially reframes post-traumatic mitophagy failure from a purely biochemical signaling deficit to a problem of pathological protein architecture, possibly underwritten by a failure of import mechanisms or aberrant degradation mechanisms [66–68]. In untreated animals, these dense TOM20 aggregates were spatially segregated from PINK1, potentially representing a physical barrier that prevents the initial docking step required to identify a damaged mitochondrion for clearance [42]. Without this initial recognition, the entire downstream rescue and removal cascade is arrested. The therapeutic action of TNS is therefore architectural; it does not simply amplify a biochemical signal but has the distinct potential to actively preserve the physical geometry of the mitochondrial surface required for that signal to function. By maintaining the diffuse, physiological distribution of TOM20 [65], TNS likely ensures that the docking sites for PINK1 remain accessible and operational [69,70]. The observation that pharmacological CGRP inhibition precipitated this pathological aggregation further provides definitive evidence that the CGRP axis directly modulates this structural preservation and is a likely causal factor behind TNS’ effects. This finding may also provide a distinct causal factor for previously observed CGRP-mediated PINK1 modulation, attributing it to more than just oxidative-stress reduction [71]. This mechanistic link, from a nerve signal, to a specific neuropeptide, to the physical arrangement of proteins on an organelle, adds a new dimension to the understanding of neuromodulation, demonstrating its capacity to exert control at a fundamental level of cellular organization.

Our study demonstrates that preserving PINK1-associated signaling through the maintenance of TOM20 integrity is linked to the protection of mitochondrial membrane integrity. Following TBI, untreated brains show a progressive loss of mitochondrial cytochrome c, spreading from the injury core outward into the penumbra and contralateral cortex. This indicates that mitochondrial membrane permeabilization is a progressive process that destabilizes surrounding and remote brain tissue. The persistence of this leakage through 72 hours corresponds with the timeline of severe mitochondrial failure and subsequent breakdown of the neuronal cytoskeleton (MAP2). TNS prevented this mitochondrial decline across all three cortical regions. In the injury core, TNS preserved mitochondrial cytochrome c despite severe primary tissue damage. In the peri-infarct penumbra, this protection lasted through the entire 72-hour period and was accompanied by the preservation of MAP2 cytoskeletal structure. The stabilization of the outer membrane’s quality control machinery, such as by the TNS-mediated preservation of the PINK1-TOM20 axis, is a key mechanism for maintaining overall mitochondrial structural integrity [72–74]. By ensuring this system remains functional, TNS exhibits the capacity to allow the cell to manage mitochondrial stress efficiently before it escalates to membrane failure. This is coupled with the robust preservation of the mitochondrial antioxidant enzyme SOD2. Together, these coordinated actions stabilize the outer mitochondrial membrane and prevent the release of cytochrome c [75]. It is plausible that by preventing the activation of the cell-death program, TNS allows neurons to survive and maintain their dendritic connections [56,57,76]. This extensive organelle preservation is a core mechanism explaining why TNS-treated animals showed a 33% reduction in brain injury volume, an 80% reduction in movement and balance deficits, and did not develop post-traumatic pain hypersensitivity [77,78]. Furthermore, this stabilized mitochondrial environment also protected the cell’s antioxidant system, NRF2 [79,80]. By preventing the initial mitochondrial failure, TNS stopped the burst of free radicals (lowering nitrotyrosine) and safely preserved nuclear NRF2 and its protective enzyme SRX1.

Our findings suggest CGRP as a critical mediator linking trigeminal neuromodulation to mitochondrial preservation after TBI. A key insight from this study is the disparity between the brain’s endogenous CGRP response to injury and the therapeutically effective response induced by TNS. The natural response is spatiotemporally dysregulated; while overall tissue CGRP levels rise, this increase does not translate into effective signaling within vulnerable neurons during the hyper-acute phase [30,43]. This results in a state where intracellular neuronal CGRP is depleted precisely when the secondary injury cascade is accelerating. TNS fundamentally corrects this pathological deficit. As a non-invasive intervention, it can impose a state of therapeutic control, driving a rapid, sustained, and neuron-centric CGRP surge that peaks within hours, not days. This contrasts with previously reported unsustained increases in CGRP following TBI [81–83], illustrating the distinction between a controlled therapeutic signal and an uncontrolled endogenous response. The necessity of this rapid CGRP signal was demonstrated by our loss-of-function experiments. Pharmacological CGRP inhibition provided a distinct exacerbation of mitochondrial structural failure and neuronal death, resulting in far more severe outcomes than those seen in untreated TBI. This experiment confirms that early CGRP availability is not an associated biomarker but is an indispensable upstream requirement for maintaining mitochondrial integrity. It is this temporal precision that is most consequential in the peri-infarct penumbra, where TNS-mediated CGRP delivery precedes the progression of dendritic decay and secondary injury expansion, confirming that the timing of the signal modulation is as important as upregulation of the signal itself. This distinction underscores the unique therapeutic advantage of TNS: it provides an external, non-pharmacological mechanism to precisely regulate the timing and localization of a critical endogenous neuropeptide, effectively overcoming a fundamental limitation of the brain’s natural injury response.

Although this study provides compelling evidence that TNS preserves mitochondrial quality control, several limitations should be noted. First, this study focused on the upstream events governing mitochondrial fate rather than directly measuring completed mitophagy flux [84,85]. Our findings indicate the primary neuroprotective mechanism is upstream mitochondrial preservation, preventing pathological TOM20 aggregation and stabilizing PINK1, which likely obviates the need for large-scale organelle degradation. While flux assays are a valuable future direction, our work identifies the decisive therapeutic action of TNS as this early structural preservation. Second, this work intimates a critical role for CGRP signaling in mediating the effects of TNS on mitochondria. Future studies using genetic models or more specific pharmacological approaches are now warranted to delineate the precise molecular interactions between CGRP, NRF2, and the PINK1/Parkin machinery. Third, we identify a novel pathological feature of TBI: the apparent aggregation of TOM20. Confirming the ultrastructure of these aggregates with advanced imaging techniques, such as cryo-electron tomography [86] or expansion microscopy [87], represents an exciting avenue to explore a new dimension of TBI pathology, and the effect which TNS has upon it. Fourth, this study focused on the acute and sub-chronic phases of TBI up to 72 hours in a rodent model. While the early reduction in injury volume and recovery in motor function are highly promising, future studies are required to confirm whether these mitochondrial benefits lead to long-term tissue remodeling and permanent behavioral recovery. Lastly, our investigation focused primarily on injured neurons; however, non-neuronal cells like microglia and astrocytes also play key roles in TBI and express CGRP receptors [30,88], meaning their mitochondrial responses to TNS deserve future investigation.

## 5. Conclusion

This study demonstrates that TNS preserves neuronal survival after TBI likely by resolving a critical structural failure at the heart of the mitochondrial quality control system. We identify this failure as the pathological aggregation of the outer mitochondrial membrane protein TOM20, which presents a likely physical obstruction to the PINK1/Parkin pathway. Crucially, our findings indicate that the key determinant of therapeutic success is not merely the presence of a protective signal, but its precise temporal delivery. TNS succeeds where the brain’s endogenous CGRP response fails because it provides this signal within the critical therapeutic window, thereby stabilizing mitochondrial architecture before cell death pathways become irreversible. By establishing the mechanistic link between trigeminal nerve stimulation and the preservation of core mitochondrial machinery, this research provides a clear and translatable strategy for mitigating the progression of post-traumatic neurodegeneration.

## Authors’ Contributions

CL conceptualized the study, administered the project, acquired the funding, provided the resources, and wrote the original draft of the manuscript. CL, KP and SW performed experiments, data collection, and data analysis. JK, KR, BT and LB contributed to the discussion and interpretation. CL, KP, JK, AD, JU, DL, LB and PW reviewed and edited the manuscript. All authors read and approved the final report and make the decision to submit for publication.

## Data Availability Statement

The data and materials supporting the findings of this study are available upon request.

## Abbreviations

A-NSS: adjusted neurological severity score
ANOVA: analysis of variance
AU: arbitrary units
BCA: bicinchoninic acid
BSA: bovine serum albumin
CCI: controlled cortical impact
CGRP: calcitonin gene-related peptide
ECL: enhanced chemiluminescence
H&E: hematoxylin and eosin
HRP: horseradish peroxidase
HSD: honestly significant difference
IV: intravenous
MAP2: microtubule-associated protein 2
NRF2: nuclear factor erythroid 2-related factor 2
OCT: optimal cutting temperature
p-NRF2: phosphorylated NRF2
PBS: phosphate-buffered saline
PFA: paraformaldehyde
PINK1: PTEN-induced kinase 1
PVDF: polyvinylidene difluoride
RIPA: radioimmunoprecipitation assay
ROI: region of interest
ROS: reactive oxygen species
SD: standard deviation
SOD2: Superoxide Dismutase 2
SRX1: Sulfiredoxin-1
TBI: traumatic brain injury
TBST: tris-buffered saline with Tween-20
TNS: trigeminal nerve stimulation
TOM20: translocase of outer mitochondrial membrane 20
VDAC1: voltage-dependent anion channel 1

## References

1. Freire, M.A.M.; Rocha, G.S.; Bittencourt, L.O.; Falcao, D.; Lima, R.R.; Cavalcanti, J.R.L.P. Cellular and Molecular Pathophysiology of Traumatic Brain Injury: What Have We Learned So Far? Biology (Basel) 2023, 12, 1139, doi:10.3390/biology12081139.

2. Rauchman, S.H.; Zubair, A.; Jacob, B.; Rauchman, D.; Pinkhasov, A.; Placantonakis, D.G.; Reiss, A.B. Traumatic Brain Injury: Mechanisms, Manifestations, and Visual Sequelae. Front Neurosci 2023, 17, 1090672, doi:10.3389/fnins.2023.1090672.

3. Maas, A.I.R.; Menon, D.K.; Manley, G.T.; Abrams, M.; Åkerlund, C.; Andelic, N.; Aries, M.; Bashford, T.; Bell, M.J.; Bodien, Y.G.;, et al. Traumatic Brain Injury: Progress and Challenges in Prevention, Clinical Care, and Research. Lancet Neurol 2022, 21, 1004– 1060, doi:10.1016/S1474-4422(22)00309-X.

4. Bahuguna, D.; Amulya, E.; Arya, S.; Loharkar, S.; Vambhurkar, G.; Bhattacharjee, S.; Chandanapalli, D.K.; Madan, J.; Srivastava, S. Unlocking Therapeutic Potential in Traumatic Brain Injury: Exploring Microenvironmental Targets, Signaling Pathways and Translational Hurdles. Inflammopharmacology 2025, 33, 5113–5144, doi:10.1007/s10787-025-01923-7.

5. Pryzmont, M.; Kosciuczuk, U.; Maciejczyk, M. Biomarkers of Traumatic Brain Injury: Narrative Review and Future Prospects in Neurointensive Care. Front Med (Lausanne) 2025, 12, 1539159, doi:10.3389/fmed.2025.1539159.

6. Katramadou, A.; Bender, E.S.; Kanakis, D. From Traumatic Brain Injury to Alzheimer’s Disease: Multilevel Biomechanical, Neurovascular, and Molecular Mechanisms with Emerging Therapeutic Directions. Int J Mol Sci 2026, 27, 1570, doi:10.3390/ijms27031570.

7. Olatona, O.A.A.; Sterben, S.P.; Kansakar, S.B.S.; Symes, A.J.J.; Liaudanskaya, V. Mitochondria: The Hidden Engines of Traumatic Brain Injury-Driven Neurodegeneration. Front. Cell. Neurosci. 2025, 19, doi:10.3389/fncel.2025.1570596.

8. Cheng, G.; Kong, R.; Zhang, L.; Zhang, J. Mitochondria in Traumatic Brain Injury and Mitochondrial-Targeted Multipotential Therapeutic Strategies. Br J Pharmacol 2012, 167, 699–719, doi:10.1111/j.1476-5381.2012.02025.x.

9. Tabassum, S.; Wu, S.; Lee, C.-H.; Yang, B.S.K.; Gusdon, A.M.; Choi, H.A.; Ren, X.S. Mitochondrial-Targeted Therapies in Traumatic Brain Injury: From Bench to Bedside. Neurotherapeutics 2025, 22, e00515, doi:10.1016/j.neurot.2024.e00515.

10. Markaki, M.; Tsagkari, D.; Tavernarakis, N. Mitophagy and Long-Term Neuronal Homeostasis. J Cell Sci 2023, 136, jcs260638, doi:10.1242/jcs.260638.

11. Yang, J.; Li, J.; Hou, X.; Zheng, Y.; Zhao, Z.; Zhou, T.; Jing, T.; Kong, J.; Zhang, G.; Guo, Y. Targeting Mitophagy for Neuroprotection: Mechanisms and Therapeutic Opportunities. NPJ Aging 2026, doi:10.1038/s41514-026-00424-3.

12. Zhu, M.; Huang, X.; Shan, H.; Zhang, M. Mitophagy in Traumatic Brain Injury: A New Target for Therapeutic Intervention. Oxid Med Cell Longev 2022, 2022, 4906434, doi:10.1155/2022/4906434.

13. Luan, Y.; Jiang, L.; Luan, Y.; Xie, Y.; Yang, Y.; Ren, K.-D. Mitophagy and Traumatic Brain Injury: Regulatory Mechanisms and Therapeutic Potentials. Oxidative Medicine and Cellular Longevity 2023, 2023, 1649842, doi:10.1155/2023/1649842.

14. Himic, V.; Tchantchaleishvili, N.; Netliukh, A.; Chibbaro, S.; Syrmos, N.; Ligarotti, G.K.I.; Prisco, L.; Ganau, M. Mitochondrial Dysfunction in Traumatic Brain Injury and Its Theranostic Implications. Biomolecules 2026, 16, 762, doi:10.3390/biom16060762.

15. Hammond, T.R.; Gadea, A.; Dupree, J.; Kerninon, C.; Nait-Oumesmar, B.; Aguirre, A.; Gallo, V. Astrocyte-Derived Endothelin-1 Inhibits Remyelination through Notch Activation. Neuron 2014, 81, 588–602, doi:10.1016/j.neuron.2013.11.015.

16. Yin, H.; Wei, Y.; Han, Z.; Li, Z.; Liu, K.; Zhu, L. BNIP3-Mediated Mitophagy Alleviates Traumatic Brain Injury by Inhibiting Neuronal Apoptosis and Oxidative Stress. Int Immunopharmacol 2026, 168, 115793, doi:10.1016/j.intimp.2025.115793.

17. Zheng, Y.; Wu, S.; Yang, Q.; Xu, Z.; Zhang, S.; Fan, S.; Liu, C.; Li, X.; Ma, C. Trigeminal Nerve Electrical Stimulation: An Effective Arousal Treatment for Loss of Consciousness. Brain Research Bulletin 2021, 169, 81–93, doi:10.1016/j.brainresbull.2021.01.008.

18. Chiluwal, A.; Narayan, R.K.; Chaung, W.; Mehan, N.; Wang, P.; Bouton, C.E.; Golanov, E.V.; Li, C. Neuroprotective Effects of Trigeminal Nerve Stimulation in Severe Traumatic Brain Injury. Sci Rep 2017, 7, 6792, doi:10.1038/s41598-017-07219-3.

19. Yang, Q.; Zhang, S.; Xu, Z.; Liu, L.; Fan, S.; Wu, S.; Ma, C. The Effectiveness of Trigeminal Nerve Stimulation on Traumatic Brain Injury. Neuromodulation 2022, 25, 1330– 1337, doi:10.1016/j.neurom.2021.10.014.

20. Xu, J.; Wu, S.; Huo, L.; Zhang, Q.; Liu, L.; Ye, Z.; Cao, J.; Ma, H.; Shang, C.; Ma, C. Trigeminal Nerve Stimulation Restores Hippocampal Dopamine Deficiency to Promote Cognitive Recovery in Traumatic Brain Injury. Prog Neurobiol 2023, 227, 102477, doi:10.1016/j.pneurobio.2023.102477.

21. Li, C.; Shah, K.A.; Powell, K.; Wu, Y.-C.; Chaung, W.; Sonti, A.N.; White, T.G.; Doobay, M.; Yang, W.-L.; Wang, P.;, et al. CBF Oscillations Induced by Trigeminal Nerve Stimulation Protect the Pericontusional Penumbra in Traumatic Brain Injury Complicated by Hemorrhagic Shock. Sci Rep 2021, 11, 19652, doi:10.1038/s41598-021-99234-8.

22. Shah, K.A.; Sonti, A.N.; Wu, Y.-C.; Powell, K.; Doobay, M.; Narayan, R.K.; Li, C. Electrical Stimulation of the Infraorbital Nerve Induces Diving Reflex in a Dose-Controlled Manner. Annu Int Conf IEEE Eng Med Biol Soc 2020, 2020, 5208–5211, doi:10.1109/EMBC44109.2020.9176845.

23. Panneton, W.M. The Mammalian Diving Response: An Enigmatic Reflex to Preserve Life? Physiology (Bethesda) 2013, 28, 284–297, doi:10.1152/physiol.00020.2013.

24. Panneton, W.M.; Gan, Q. The Mammalian Diving Response: Inroads to Its Neural Control. Front Neurosci 2020, 14, 524, doi:10.3389/fnins.2020.00524.

25. Li, C.; Powell, K.; Wadolowski, S.; Jacob, A.; LeDoux, C.; Brenner, M.; Wang, P. The Diving Reflex Elicits Potent, Non-Electrophilic Nrf2 Activation via Trigeminal CGRP Signaling. Free Radic Biol Med 2026, 253, 238–256, doi:10.1016/j.freeradbiomed.2026.05.306.

26. Tambo, W.; Powell, K.; Wadolowski, S.; Unadkat, P.; Chang, E.H.; LeDoux, C.; Sciubba, D.; Wang, P.; Huerta, P.; Li, C. Vasoactive Neuropeptide Dysregulation: A Novel Mechanism of Microvascular Dysfunction in Vascular Cognitive Impairment. Alzheimers Dement 2025, 21, e70925, doi:10.1002/alz.70925.

27. Ngo, V.; Duennwald, M.L. Nrf2 and Oxidative Stress: A General Overview of Mechanisms and Implications in Human Disease. Antioxidants (Basel) 2022, 11, 2345, doi:10.3390/antiox11122345.

28. Esteras, N.; Abramov, A.Y. Nrf2 as a Regulator of Mitochondrial Function: Energy Metabolism and Beyond. Free Radic Biol Med 2022, 189, 136–153, doi:10.1016/j.freeradbiomed.2022.07.013.

29. Li, C.; White, T.G.; Shah, K.A.; Chaung, W.; Powell, K.; Wang, P.; Woo, H.H.; Narayan, R.K. Percutaneous Trigeminal Nerve Stimulation Induces Cerebral Vasodilation in a Dose-Dependent Manner. Neurosurgery 2021, 88, E529–E536, doi:10.1093/neuros/nyab053.

30. Lin, K.; Stiles, J.; Tambo, W.; Ajmal, E.; Piao, Q.; Powell, K.; Li, C. Bimodal Functions of Calcitonin Gene-Related Peptide in the Brain. Life Sciences 2024, 359, 123177, doi:10.1016/j.lfs.2024.123177.

31. Xiong, J.; Wang, Z.; Bai, J.; Cheng, K.; Liu, Q.; Ni, J. Calcitonin Gene-Related Peptide: A Potential Protective Agent in Cerebral Ischemia–Reperfusion Injury. Front Neurosci 2023, 17, 1184766, doi:10.3389/fnins.2023.1184766.

32. Kee, Z.; Kodji, X.; Brain, S.D. The Role of Calcitonin Gene Related Peptide (CGRP) in Neurogenic Vasodilation and Its Cardioprotective Effects. Front Physiol 2018, 9, 1249, doi:10.3389/fphys.2018.01249.

33. Gumeni, S.; Papanagnou, E.-D.; Manola, M.S.; Trougakos, I.P. Nrf2 Activation Induces Mitophagy and Reverses Parkin/Pink1 Knock down-Mediated Neuronal and Muscle Degeneration Phenotypes. Cell Death Dis 2021, 12, 671, doi:10.1038/s41419-021-03952-w.

34. Gureev, A.P.; Sadovnikova, I.S.; Starkov, N.N.; Starkov, A.A.; Popov, V.N. P62-Nrf2-P62 Mitophagy Regulatory Loop as a Target for Preventive Therapy of Neurodegenerative Diseases. Brain Sci 2020, 10, 847, doi:10.3390/brainsci10110847.

35. Ryoo, I.-G.; Kwak, M.-K. Regulatory Crosstalk between the Oxidative Stress-Related Transcription Factor Nfe2l2/Nrf2 and Mitochondria. Toxicol Appl Pharmacol 2018, 359, 24– 33, doi:10.1016/j.taap.2018.09.014.

36. Bingol, B.; Sheng, M. Mechanisms of Mitophagy: PINK1, Parkin, USP30 and Beyond. Free Radic Biol Med 2016, 100, 210–222, doi:10.1016/j.freeradbiomed.2016.04.015.

37. Lamade, A.M.; Anthonymuthu, T.S.; Hier, Z.E.; Gao, Y.; Kagan, V.E.; Bayır, H. Mitochondrial Damage & Lipid Signaling in Traumatic Brain Injury. Exp Neurol 2020, 329, 113307, doi:10.1016/j.expneurol.2020.113307.

38. Lin, C.; Li, N.; Chang, H.; Shen, Y.; Li, Z.; Wei, W.; Chen, H.; Lu, H.; Ji, J.; Liu, N. Dual Effects of Thyroid Hormone on Neurons and Neurogenesis in Traumatic Brain Injury. Cell Death Dis 2020, 11, 671, doi:10.1038/s41419-020-02836-9.

39. Li, Y.; Chen, W.; Feng, J. Astrocyte-Derived Exosomal GJA1-20 k Targets Pink1-Mediated Mitophagy to Attenuate Traumatic Brain Injury. Transl Stroke Res 2025, 16, 2119–2129, doi:10.1007/s12975-025-01374-w.

40. Wenzhe, L.; Boyang, X.; Yuchao, G.; Bimcle, R.; Yue, Y. Mitochondrial and ER Stress Crosstalk in TBI: Mechanistic Insights and Therapeutic Opportunities. Front Cell Neurosci 2025, 19, 1697060, doi:10.3389/fncel.2025.1697060.

41. Barodia, S.K.; Creed, R.B.; Goldberg, M.S. Parkin and PINK1 Functions in Oxidative Stress and Neurodegeneration. Brain Res Bull 2017, 133, 51–59, doi:10.1016/j.brainresbull.2016.12.004.

42. Eldeeb, M.A.; Bayne, A.N.; Fallahi, A.; Goiran, T.; MacDougall, E.J.; Soumbasis, A.; Zorca, C.E.; Tabah, J.-J.; Thomas, R.A.; Karpilovsky, N.;, et al. Tom20 Gates PINK1 Activity and Mediates Its Tethering of the TOM and TIM23 Translocases upon Mitochondrial Stress. Proc Natl Acad Sci U S A 121, e2313540121, doi:10.1073/pnas.2313540121.

43. Li, C.; Ajmal, E.; Alok, K.; Powell, K.; Wadolowski, S.; Tambo, W.; Turpin, J.; Barthélemy, E.; Al-Abed, Y.; LeDoux, D. CGRP as a Potential Mediator for the Sexually Dimorphic Responses to Traumatic Brain Injury. Biol Sex Differ 2024, 15, 44, doi:10.1186/s13293-024-00619-x.

44. Lynch, D.G.; Shah, K.A.; Powell, K.; Wadolowski, S.; Tambo, W.; Strohl, J.J.; Unadkat, P.; Eidelberg, D.; Huerta, P.T.; Li, C. Neurobehavioral Impairments Predict Specific Cerebral Damage in Rat Model of Subarachnoid Hemorrhage. Transl Stroke Res 2023, doi:10.1007/s12975-023-01180-2.

45. Santos-Nogueira, E.; Redondo Castro, E.; Mancuso, R.; Navarro, X. Randall-Selitto Test: A New Approach for the Detection of Neuropathic Pain after Spinal Cord Injury. J Neurotrauma 2012, 29, 898–904, doi:10.1089/neu.2010.1700.

46. Yam, M.F.; Loh, Y.C.; Oo, C.W.; Basir, R. Overview of Neurological Mechanism of Pain Profile Used for Animal “Pain-Like” Behavioral Study with Proposed Analgesic Pathways. Int J Mol Sci 2020, 21, 4355, doi:10.3390/ijms21124355.

47. Geisert, E.E.; Johnson, H.G.; Binder, L.I. Expression of Microtubule-Associated Protein 2 by Reactive Astrocytes. Proc Natl Acad Sci U S A 1990, 87, 3967–3971, doi:10.1073/pnas.87.10.3967.

48. Matsunaga, W.; Miyata, S.; Kiyohara, T. Redistribution of MAP2 Immunoreactivity in the Neurohypophysial Astrocytes of Adult Rats during Dehydration. Brain Res 1999, 829, 7– 17, doi:10.1016/s0006-8993(99)01285-8.

49. Goemans, C.G.; Boya, P.; Skirrow, C.J.; Tolkovsky, A.M. Intra-Mitochondrial Degradation of Tim23 Curtails the Survival of Cells Rescued from Apoptosis by Caspase Inhibitors. Cell Death Differ 2008, 15, 545–554, doi:10.1038/sj.cdd.4402290.

50. Thayer, J.A.; Petersen, J.D.; Huang, X.; Gruel Budet, L.M.; Hawrot, J.; Ramos, D.M.; Sekine, S.; Li, Y.; Ward, M.E.; Narendra, D.P. A Unified Mechanism for Mitochondrial Damage Sensing in PINK1-Parkin-Mediated Mitophagy. EMBO J 2026, 45, 64–105, doi:10.1038/s44318-025-00604-z.

51. Callegari, S.; Cruz-Zaragoza, L.D.; Rehling, P. From TOM to the TIM23 Complex - Handing over of a Precursor. Biol Chem 2020, 401, 709–721, doi:10.1515/hsz-2020-0101.

52. Franco-Iborra, S.; Cuadros, T.; Parent, A.; Romero-Gimenez, J.; Vila, M.; Perier, C. Defective Mitochondrial Protein Import Contributes to Complex I-Induced Mitochondrial Dysfunction and Neurodegeneration in Parkinson’s Disease. Cell Death Dis 2018, 9, 1122, doi:10.1038/s41419-018-1154-0.

53. Yamano, K.; Youle, R.J. PINK1 Is Degraded through the N-End Rule Pathway. Autophagy 2013, 9, 1758–1769, doi:10.4161/auto.24633.

54. Shiba-Fukushima, K.; Imai, Y.; Yoshida, S.; Ishihama, Y.; Kanao, T.; Sato, S.; Hattori, N. PINK1-Mediated Phosphorylation of the Parkin Ubiquitin-like Domain Primes Mitochondrial Translocation of Parkin and Regulates Mitophagy. Sci Rep 2012, 2, 1002, doi:10.1038/srep01002.

55. Connelly, E.M.; Frankel, K.S.; Shaw, G.S. Parkin and Mitochondrial Signalling. Cell Signal 2023, 106, 110631, doi:10.1016/j.cellsig.2023.110631.

56. Eleftheriadis, T.; Pissas, G.; Liakopoulos, V.; Stefanidis, I. Cytochrome c as a Potentially Clinical Useful Marker of Mitochondrial and Cellular Damage. Front Immunol 2016, 7, 279, doi:10.3389/fimmu.2016.00279.

57. Waterhouse, N.J.; Goldstein, J.C.; von Ahsen, O.; Schuler, M.; Newmeyer, D.D.; Green, D.R. Cytochrome c Maintains Mitochondrial Transmembrane Potential and ATP Generation after Outer Mitochondrial Membrane Permeabilization during the Apoptotic Process. J Cell Biol 2001, 153, 319–328, doi:10.1083/jcb.153.2.319.

58. Campos, C.B.L.; Paim, B.A.; Cosso, R.G.; Castilho, R.F.; Rottenberg, H.; Vercesi, A.E. Method for Monitoring of Mitochondrial Cytochrome c Release during Cell Death: Immunodetection of Cytochrome c by Flow Cytometry after Selective Permeabilization of the Plasma Membrane. Cytometry A 2006, 69, 515–523, doi:10.1002/cyto.a.20273.

59. Balan, I.S.; Saladino, A.J.; Aarabi, B.; Castellani, R.J.; Wade, C.; Stein, D.M.; Eisenberg, H.M.; Chen, H.H.; Fiskum, G. Cellular Alterations in Human Traumatic Brain Injury: Changes in Mitochondrial Morphology Reflect Regional Levels of Injury Severity. J Neurotrauma 2013, 30, 367–381, doi:10.1089/neu.2012.2339.

60. Lifshitz, J.; Friberg, H.; Neumar, R.W.; Raghupathi, R.; Welsh, F.A.; Janmey, P.; Saatman, K.E.; Wieloch, T.; Grady, M.S.; McIntosh, T.K. Structural and Functional Damage Sustained by Mitochondria after Traumatic Brain Injury in the Rat: Evidence for Differentially Sensitive Populations in the Cortex and Hippocampus. J Cereb Blood Flow Metab 2003, 23, 219– 231, doi:10.1097/01.WCB.0000040581.43808.03.

61. Wu, H.-M.; Huang, S.-C.; Vespa, P.; Hovda, D.A.; Bergsneider, M. Redefining the Pericontusional Penumbra Following Traumatic Brain Injury: Evidence of Deteriorating Metabolic Derangements Based on Positron Emission Tomography. J Neurotrauma 2013, 30, 352–360, doi:10.1089/neu.2012.2610.

62. Tian, J.; Mao, Y.; Liu, D.; Li, T.; Wang, Y.; Zhu, C. Mitophagy in Brain Injuries: Mechanisms, Roles, and Therapeutic Potential. Mol Neurobiol 2025, 62, 10856–10868, doi:10.1007/s12035-025-04936-z.

63. Wu, X.; Lv, W.; Chen, H.; Fu, Y.; Fei, X.; Dou, Y.; Luo, P.; Yu, Y.; Niu, Z.; Huan, Y.;, et al. Parkin Protects against Traumatic Brain Injury through Regulating Mitochondrial Quality Control. Neurotherapeutics 2026, 23, e00846, doi:10.1016/j.neurot.2026.e00846.

64. Yano, M.; Terada, K.; Mori, M. Mitochondrial Import Receptors Tom20 and Tom22 Have Chaperone-like Activity. J Biol Chem 2004, 279, 10808–10813, doi:10.1074/jbc.M311710200.

65. Wurm, C.A.; Neumann, D.; Lauterbach, M.A.; Harke, B.; Egner, A.; Hell, S.W.; Jakobs, S. Nanoscale Distribution of Mitochondrial Import Receptor Tom20 Is Adjusted to Cellular Conditions and Exhibits an Inner-Cellular Gradient. Proc Natl Acad Sci U S A 2011, 108, 13546–13551, doi:10.1073/pnas.1107553108.

66. Nowicka, U.; Chroscicki, P.; Stroobants, K.; Sladowska, M.; Turek, M.; Uszczynska-Ratajczak, B.; Kundra, R.; Goral, T.; Perni, M.; Dobson, C.M.;, et al. Cytosolic Aggregation of Mitochondrial Proteins Disrupts Cellular Homeostasis by Stimulating the Aggregation of Other Proteins. Elife 2021, 10, e65484, doi:10.7554/eLife.65484.

67. Liu, Q.; Fong, B.; Yoo, S.; Unruh, J.R.; Guo, F.; Yu, Z.; Chen, J.; Si, K.; Li, R.; Zhou, C. Nascent Mitochondrial Proteins Initiate the Localized Condensation of Cytosolic Protein Aggregates on the Mitochondrial Surface. Proc Natl Acad Sci U S A 2023, 120, e2300475120, doi:10.1073/pnas.2300475120.

68. Hatori, Y.; Kanda, Y.; Nonaka, S.; Nakanishi, H.; Kitazawa, T. ATP13A2 Modifies Mitochondrial Localization of Overexpressed TOM20 to Autolysosomal Pathway. PLoS One 2022, 17, e0276823, doi:10.1371/journal.pone.0276823.

69. Borrero-Landazabal, M.A.; Linke, V.; Kadavá, T.; Li, Z.; Draczkowski, P.; Kaszuba, K.; Bertgen, L.; Serwa, R.A.; Heck, A.J.R.; Chacinska, A. A Dynamic Gateway: Uncovering the Expanded Human TOM Complex Interactome and Its Regulatory Complexity. Sci Adv 2026, 12, eaeb2995, doi:10.1126/sciadv.aeb2995.

70. Su, J.; Tian, X.; Wang, Z.; Yang, J.; Sun, S.; Sui, S.-F. Structure of the Intact Tom20 Receptor in the Human Translocase of the Outer Membrane Complex. PNAS Nexus 2024, 3, pgae269, doi:10.1093/pnasnexus/pgae269.

71. Mishra, A.; Betancourt, A.; Vidyadharan, V.A.; Blesson, C.S.; Belfort, M.; Yallampalli, C.; Chauhan, M. Calcitonin Gene-Related Peptide Protects from Soluble Fms-like Tyrosine Kinase-1-Induced Vascular Dysfunction in a Preeclampsia Mouse Model. Front Physiol 2023, 14, 1221684, doi:10.3389/fphys.2023.1221684.

72. Pitt, A.S.; Buchanan, S.K. A Biochemical and Structural Understanding of TOM Complex Interactions and Implications for Human Health and Disease. Cells 2021, 10, 1164, doi:10.3390/cells10051164.

73. Maruszczak, K.K.; Jung, M.; Rasool, S.; Trempe, J.-F.; Rapaport, D. The Role of the Individual TOM Subunits in the Association of PINK1 with Depolarized Mitochondria. J Mol Med (Berl) 2022, 100, 747–762, doi:10.1007/s00109-022-02191-6.

74. Collier, C.L.; Ruedi, C.; Thorne, N.J.; Tumbarello, D.A. PINK1 Loss of Function Selectively Alters the Mitochondrial-Derived Vesicle Pathway. FASEB Bioadv 2025, 7, e70030, doi:10.1096/fba.2024-00200.

75. Zhang, Z.; Zhang, M.; Jin, H.; Lv, S.; Li, Y.; Li, Y. Mitochondrial Quality Control and Cell Death. Int J Mol Sci 2025, 26, 11084, doi:10.3390/ijms262211084.

76. Zhou, Z.; Arroum, T.; Luo, X.; Kang, R.; Lee, Y.J.; Tang, D.; Hüttemann, M.; Song, X. Diverse Functions of Cytochrome c in Cell Death and Disease. Cell Death Differ 2024, 31, 387–404, doi:10.1038/s41418-024-01284-8.

77. Benson, C.A.; King, J.F.; Reimer, M.L.; Kauer, S.D.; Waxman, S.G.; Tan, A.M. Dendritic Spines and Pain Memory. Neuroscientist 2024, 30, 294–314, doi:10.1177/10738584221138251.

78. Kauer, S.D.; Effraim, P.R.; Bangalore, L.; Waxman, S.G.; Tan, A.M. Dendritic Spine Dysgenesis in Spinal Cord Injury: A Structural Contributor to Pain and Spasticity. Exp Neurol 2026, 399, 115679, doi:10.1016/j.expneurol.2026.115679.

79. Saso, L.; Ates, I.; Tunc, R.; Yilmaz, B.; Gallorini, M.; Carradori, S.; Suzen, S. Modulation of Nrf2 and Mitochondrial Function: Pharmacological Implications. Pharmaceuticals (Basel) 2025, 18, 1698, doi:10.3390/ph18111698.

80. Cvetko, F.; Caldwell, S.T.; Higgins, M.; Suzuki, T.; Yamamoto, M.; Prag, H.A.; Hartley, R.C.; Dinkova-Kostova, A.T.; Murphy, M.P. Nrf2 Is Activated by Disruption of Mitochondrial Thiol Homeostasis but Not by Enhanced Mitochondrial Superoxide Production. J Biol Chem 2021, 296, 100169, doi:10.1074/jbc.RA120.016551.

81. Chen, L.-X.; Zhang, W.-F.; Wang, M.; Jia, P.-F. Relationship of Calcitonin Gene-Related Peptide with Disease Progression and Prognosis of Patients with Severe Traumatic Brain Injury. Neural Regen Res 2018, 13, 1782–1786, doi:10.4103/1673-5374.238619.

82. Hang, C.-H.; Shi, J.-X.; Li, J.-S.; Wu, W.; Li, W.-Q.; Yin, H.-X. Levels of Vasoactive Intestinal Peptide, Cholecystokinin and Calcitonin Gene-Related Peptide in Plasma and Jejunum of Rats Following Traumatic Brain Injury and Underlying Significance in Gastrointestinal Dysfunction. World J Gastroenterol 2004, 10, 875–880, doi:10.3748/wjg.v10.i6.875.

83. Naseri Alavi, S.A.; Habibi, M.A.; Rezakhah, A.; Naseri Alavi, S.H.; Heydarian, P.; Torkan, J.S.M.; Jung, G.; Keymakh, M.; Kobets, A.J. Prognostic Serum Calcitonin Gene-Related Peptide Level Value in Patients Following Traumatic Brain Injury. Asian J Neurosurg 2025, 20, 285–290, doi:10.1055/s-0044-1801783.

84. Singh, K.; Berg, T. Extracellular Flux Assays for the Measurement of Glycolysis and Mitochondrial Respiration in Brain Cancer Cells. Methods Mol Biol 2025, 2944, 73–109, doi:10.1007/978-1-0716-4654-0_7.

85. Sharma, E.; Fotooh Abadi, L.; Kombe Kombe, J.A.; Kandala, M.; Parker, J.; Winicki, N.; Kelesidis, T. Overview of Methods That Determine Mitochondrial Function in Human Disease. Metabolism 2025, 170, 156300, doi:10.1016/j.metabol.2025.156300.

86. Agip, A.-N.A.; Ornelas, P.; Yang, T.-J.; Uboldi, E.; Häder, S.; McDowell, M.A.; Kühlbrandt, W. Structures of Chaetomium Thermophilum TOM Complexes with Bound Preproteins. Proc Natl Acad Sci U S A 2025, 122, e2507279122, doi:10.1073/pnas.2507279122.

87. Kunz, T.C.; Götz, R.; Gao, S.; Sauer, M.; Kozjak-Pavlovic, V. Using Expansion Microscopy to Visualize and Characterize the Morphology of Mitochondrial Cristae. Front Cell Dev Biol 2020, 8, 617, doi:10.3389/fcell.2020.00617.

88. Karve, I.P.; Taylor, J.M.; Crack, P.J. The Contribution of Astrocytes and Microglia to Traumatic Brain Injury. Br J Pharmacol 2016, 173, 692–702, doi:10.1111/bph.13125.

