## Supplemental Materials for "Trigeminal Nerve Stimulation Attenuates Traumatic Brain Injury–Induced Neuronal Damage via CGRP-PINK1 Mitophagy Axis"

**This word file includes:**

- **Supplementary Tables 1-2**
- **Supplementary Figures 1-4**

**Supplementary Table 1. Product information of commercial kits**

| **Product Name** | **Vendor** | **SKU** |
| --- | --- | --- |
| NE-PER™ Nuclear and Cytoplasmic Extraction Reagents | Thermo Scientific | 78835 |
| GSH/GSSG Ratio Detection Assay Kit (Fluorometric - Green) | Abcam | ab138881 |
| Lipid Peroxidation (MDA) Assay Kit (Colorimetric/Fluorometric) | Abcam | ab118970 |
| Mitochondria | Abcam | ab110168 |

**Supplementary Table 2. The antibodies used in this study**

| **Antibodies** | **Source** | **SKU** | **Host** | **Application and dilution** |
| --- | --- | --- | --- | --- |
| CGRP | Santa Cruz Biotechnology | sc-57053 | ms | Western blotting, 1:100 in 5% milk in TBST |
| CGRP | Abcam | ab81887 | ms | Immunofluorescence, 1:50 in 1% BSA in TBST |
| NRF2 | Proteintech | 16396-1-AP | rb | Western blotting, 1:1000 in 5% BSA in TBST |
| Phosphorylated NRF2, s40 | Abcam | AB76026 | rb | Immunofluorescence, 1:500 in 1% BSA in TBST |
| LC3II | Proteintech | 18725-1-AP | Rb | Western blotting, 1:1000 in 5% milk in TBST |
| SOD2 | Proteintech | 24127-1-AP | rb | Western blotting, 1:10000 in 5% milk in TBST |
| Cytochrome c | Proteintech | 66264-1-Ig | ms | Western blotting, 1:5000 in 5% milk in TBST  Immunofluorescence, 1:500 in 1% BSA in TBST |
| PINK1 | Proteintech | 23274-1-AP | rb | Western blotting, 1:1000 in 5% milk in TBST  Immunofluorescence, 1:500 in 1% BSA in TBST |
| Parkin | Proteintech | 66674-4-Ig | Ms | Western blotting, 1:5000 in 5% milk in TBST |
| Nitrotyrosine | Santa Cruz Biotechnology | Sc-101358 | ms | Immunofluorescence, 1:100 in 1% BSA in TBST |
| SRX1 | Proteintech | 14273-1-AP | rb | Western blotting, 1:3000 in 5% milk in TBST |
| 4-HNE | Abcam | ab48506 | N/A | Immunofluorescence, 1:25 in 1% BSA in TBST |
| TOM20 | Santa Cruz Biotechnology | SC-136211 | ms | Immunofluorescence, 1:100 in 1% BSA in TBST |
| NeuN | Abcam | Ab104224 | ms | Immunofluorescence, 1:500 in 1% BSA in TBST |
| NeuN | Abcam | Ab177487 | rb | Immunofluorescence, 1:100 in 1% BSA in TBST |
| MAP2 | Abcam | Ab32454 | ms | Western blotting, 1:1000 in 5% milk in TBST  Immunofluorescence, 1:500 in 1% BSA in TBST |
| β-actin | Sigma | A5441-.2ML | ms | Western blotting, 1:25000 in 5% milk in TBST |
| VDAC1 | Proteintech | 66345-1-Ig | ms | Western blotting, 1:5000 in 5% milk in TBST |
| Lamin B1 | Proteintech | 66095-1-IG | ms | Western blotting, 1:5000 in 5% milk in TBST |
| DAPI Solution | Thermo Fisher Scientific | 62248 | N/A | Immunofluorescence, 1:2000 |

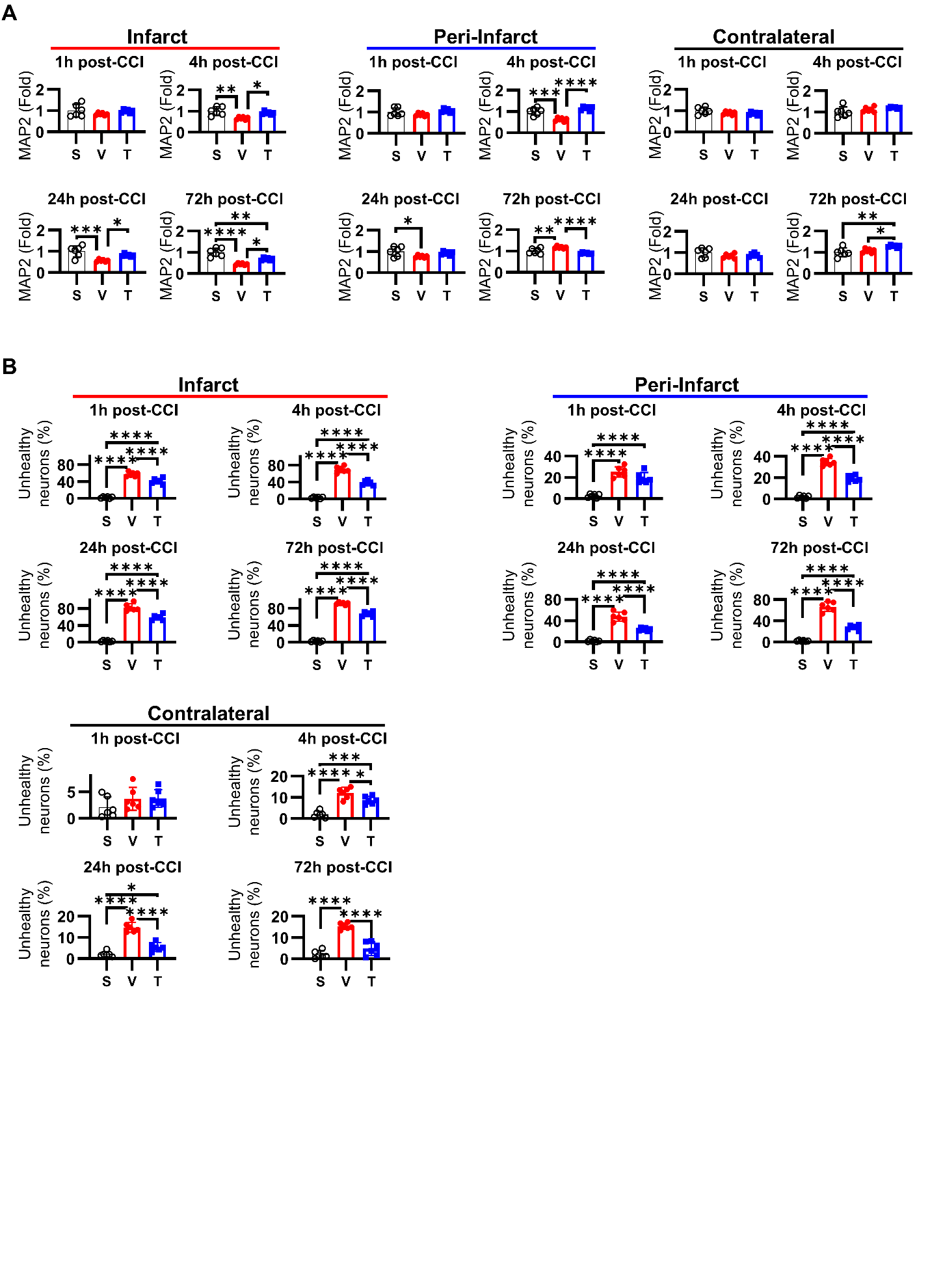

**Supplementary Figure 1. TNS mitigates post-traumatic cytoskeletal degradation and neuronal degeneration. (A)** Quantification of Western blot data showing the time course of MAP2 protein expression at 1, 4, 24, and 72 hours post-CCI in the infarct core, peri-infarct penumbra, and contralateral cortex. **(B)** Quantification of unhealthy neurons, identified by condensed or pyknotic nuclei, at the same time points and across the same three cortical regions. TNS treatment significantly reduced the percentage of unhealthy neurons compared to vehicle-treated controls, particularly in the peri-infarct penumbra.

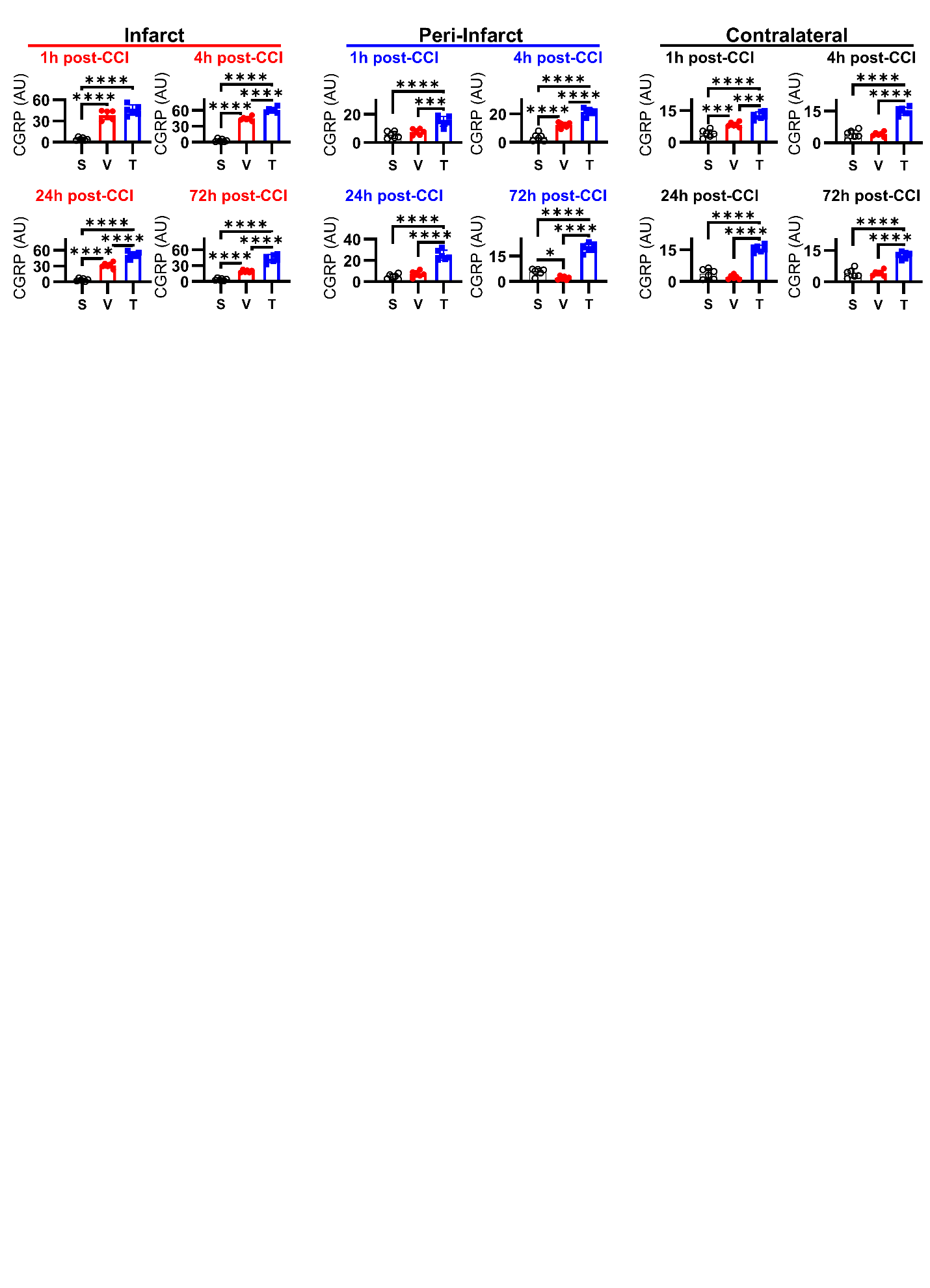

**Supplementary Figure 2. TNS drives an early and sustained upregulation of neuronal CGRP after TBI.** Quantification of neuronal CGRP expression from immunofluorescence imaging at 1, 4, 24, and 72 hours post-CCI. Data are shown for the infarct core, peri-infarct penumbra, and contralateral cortex. TNS treatment induced a rapid and sustained elevation of CGRP within neurons, particularly in the infarct and peri-infarct regions, preventing the transient and ultimately depleted response seen in vehicle-treated animals.

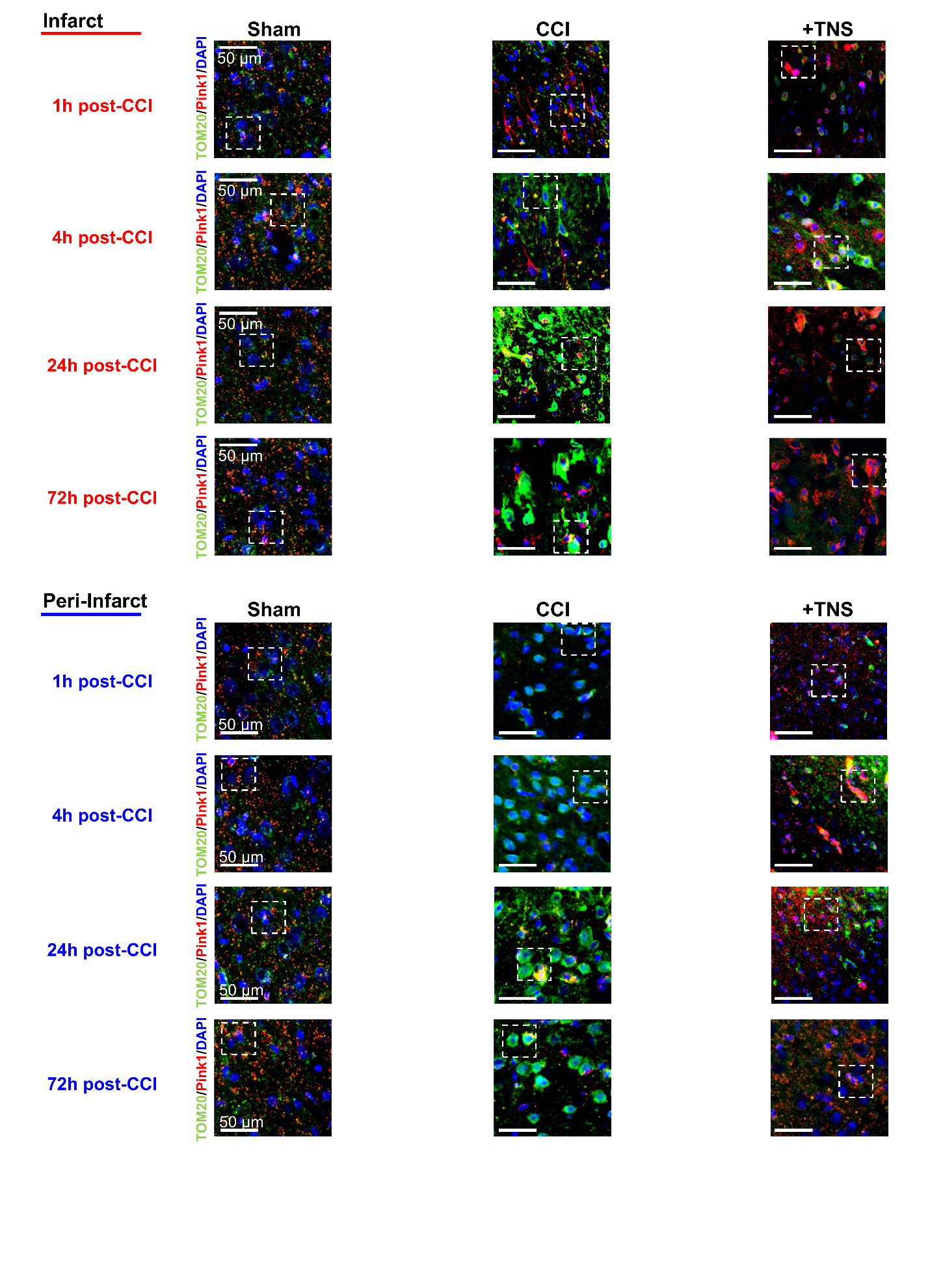

**Supplementary Figure 3. Large-field views showing the spatiotemporal dynamics of TOM20 and PINK1 following TBI.** Large region-of-interest (ROI) immunofluorescence images showing the distribution of the outer mitochondrial membrane protein TOM20 (green) and the mitophagy sensor PINK1 (red) at 1, 4, 24, and 72 hours post-CCI in both the infarct core and peri-infarct cortex. These images provide broader context for the high-magnification views shown in Figure 5, demonstrating the widespread nature of the pathological TOM20 aggregation in vehicle-treated animals and its resolution with TNS treatment over time.

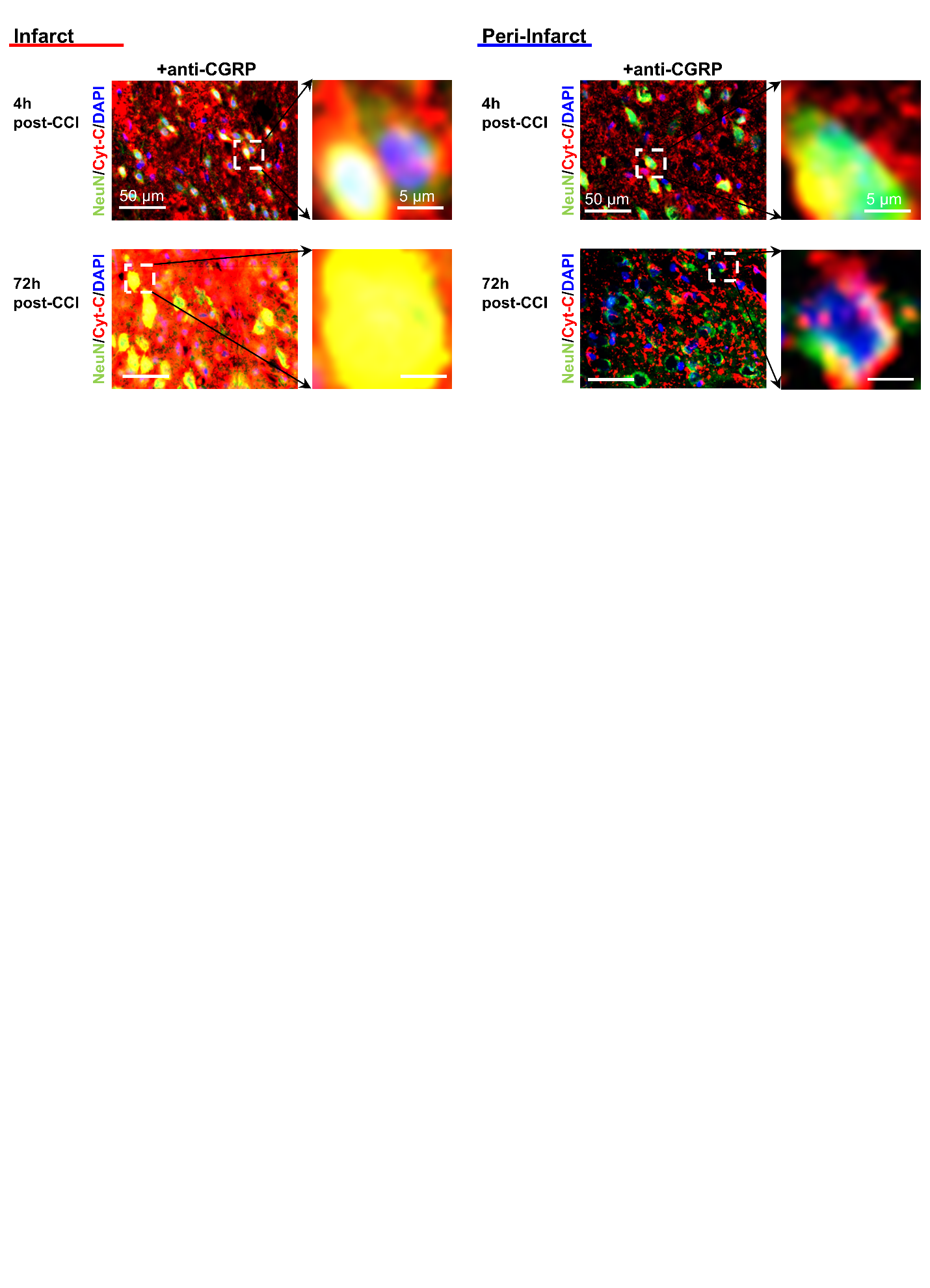

**Supplementary Figure 4. CGRP inhibition exacerbates post-traumatic neuronal cytochrome c (Cyt-C) leakage.** Representative immunofluorescence images showing the effect of pharmacological CGRP receptor inhibition on neuronal cytochrome c leakage at 4 and 72 hours post-CCI in both the infarct core and peri-infarct cortex. The images demonstrate that blocking CGRP signaling significantly worsens the cytoplasmic accumulation of cytochrome c compared to vehicle-treated TBI animals, confirming that endogenous CGRP is required to maintain mitochondrial outer membrane integrity and prevent the release of this pro-apoptotic factor. This provides the direct visual evidence supporting the conclusion in Figure 7C.
